# Evidence for the Type IV Pili Retraction Motor PilT as a Component of the Surface Sensing System in *Pseudomonas aeruginosa*

**DOI:** 10.1101/2023.05.02.539127

**Authors:** C.J. Geiger, G.A. O’Toole

## Abstract

Biofilm formation begins when bacteria contacting a surface induce cellular changes to become better adapted for surface growth. One of the first changes to occur for *Pseudomonas aeruginosa* after surface contact is an increase in the nucleotide second messenger 3’,5’-cyclic adenosine monophosphate (cAMP). It has been demonstrated that this increase in intracellular cAMP is dependent on functional Type IV pili (T4P) relaying a signal to the Pil-Chp system, but the mechanism by which this signal is transduced remains poorly understood. Here, we investigate the role of the Type IV pili retraction motor PilT in sensing a surface and relaying that signal to cAMP production. We show that mutations affecting the structure of PilT and in particular ATPase activity of this motor protein, reduce surface-dependent cAMP production. We identify a novel interaction between PilT and PilJ, a member of the Pil-Chp system, and propose a new model whereby *P. aeruginosa* uses its retraction motor to sense a surface and to relay that signal via PilJ to increased production of cAMP. We discuss these findings in light of current TFP-dependent surface sensing models for *P. aeruginosa*.

**Importance:** T4P are cellular appendages that allow *P. aeruginosa* to sense a surface leading to the production of cAMP. This second messenger not only activates virulence pathways but leads to further surface adaptation and irreversible attachment of cells. Here, we demonstrate the importance of the retraction motor PilT in surface sensing. We also present a new surface sensing model in *P. aeruginosa* whereby the T4P retraction motor PilT senses and transmits the surface signal, likely via its ATPase domain and interaction with PilJ, to mediate production of the second messenger cAMP.

## Introduction

Biofilm formation is initiated when free swimming, planktonic cells contact a surface. This contact serves as a signal that must be transmitted across the cell envelope into the cytoplasm to initiate appropriate physiological changes to adapt to the biofilm mode of growth (1). For many bacteria this initial surface contact is mediated through motility appendages such as type IV pili (T4P) or flagella (2–8). Contact between these appendages and the surface creates forces that are not normally present in planktonic environments and can serve as a “surface signal” to the microbe (9).

Early work in *Vibrio parahaemolyticus* demonstrated that the signals encountered during surface contact could be mimicked by increasing the load on the flagellum either through changes in viscosity of the medium or by addition of antibodies specific to the flagellum (10, 11). Recent work in *Caulobacter crescentus* demonstrated that holdfast formation and DNA replication, which normally occurs during surface contact, could be stimulated by increasing the load on Tad pili during retraction. Furthermore, the baseline number of cells with a holdfast without prior pili obstruction was higher in mutants that were unable to rotate their flagellum (2). Others have demonstrated that the flagellar motor itself is able to sense surface contact to trigger c-di-GMP production leading to holdfast synthesis (12). Together, these data indicate that bacteria use their surface appendages to help sense surface engagement and indicate that impeding the motion (i.e., retraction and/or rotation) of these appendages might serve as the proximal signal for surface engagement.

*Pseudomonas aeruginosa*, like other organisms, utilizes T4P as well as its polar flagellum to sense and traverse surfaces (3, 13–16). One of the first changes to occur for many organisms upon surface contact is an increase in the second messenger cyclic-di-GMP (cdG) (17). In *P. aeruginosa* PA14, this initial increase in cdG is produced by the diguanylate cyclase SadC and recent work from our lab and others has shown that SadC activity is regulated by both components of the flagellum and the T4P (3, 18). Prior to an increase in cdG, *P. aeruginosa* PA14 increases levels of another second messenger, 3’,5’-cyclic adenosine monophosphate (cAMP) (15).

The surface-dependent increase of cAMP by *P. aeruginosa* PA14 depends on functional T4P, the Pil-Chp chemotaxis-like system, and the adenylate cyclase CyaB, and to a lesser extent, the adenylate cyclase CyaA. The methyl-accepting chemotaxis protein (MCP) PilJ relays a signal to the kinase ChpA (19). Activation of the system causes ChpA to phosphorylate the response regulator PilG; phosphorylation of PilG as well as FimV and FimL are required to then activate the adenylate cyclases CyaAB to produce cAMP from ATP (20–22). The cAMP binding protein Vfr then binds cAMP and activates genes necessary for further surface adaptation, as well as for virulence (23, 24).

Recent work by Yarrington, Limoli and colleagues shows that the PilJ likely detects phenyl soluble modulins via its periplasmic domain as a ligand to trigger signaling, a finding that strongly suggests that PilJ can function like a classic MCP (25). Others have recently uncovered the function of PilG and PilH in twitching motility and surface adaptation (26, 27). In contrast, how surface engagement by T4P triggers cAMP signaling in a PilJ-dependent manner is still an open question. A previous study showed that the ligand binding domain (LBD) of PilJ is not required for surface-dependent cAMP production, although the extent of cAMP induction is significantly reduced relative to the WT (28). One model to explain TFP-mediated surface signaling includes interactions between PilA-PilJ via a “mechanosensing” signal; we recently reported data at odds with this model (29).

*P. aeruginosa* utilizes the T4P as a cellular grappling hook that pulls the cells along a surface through rounds of pilus extension, surface binding and pilus retraction (14). Functional pili are also required for sensitivity to infection by the phage DMS3 (30). Extension and retraction are powered by three hexameric ATPases: PilB, PilT, and PilU (31–33). In a recent study from our group, we found that surface pili and surface engagement are required for surface signaling, consistent with previous studies (15, 19, 23, 34, 35). Furthermore, we showed that the ability to retract pili with only enough force to allow phage infection was necessary for surface-dependent, cAMP production. That is, the force required for twitching motility was not necessary for cAMP signaling (29). Since phage susceptibility and the cAMP response both require the retraction motor PilT, we reasoned that PilT may be involved in surface-mediated signal transduction in addition to its role in pilus retraction.

To explore the potential role of PilT in surface signaling, we began by characterizing the effect of different PilT mutations on surface dependent cAMP production during biofilm formation. We found that mutations in PilT affecting ATP binding and hydrolysis affected cAMP production. A Bacterial Adenylate Cyclase Two Hybrid (B2H) screen revealed a novel interaction between PilT and PilJ. We screened for mutant PilT alleles with altered interaction with PilJ, but which retained the ability to perform twitching motility. We report here a strong relationship between the extent of PilT-PilJ interaction for PilT mutants that are defective in ATPase activity and the magnitude of cAMP signaling. We also find a strong relationship between the twitching motility zone size and the extent of cAMP production. We also identify a mutation in PilT that disrupts its interaction with PilJ in a B2H assay in *E. coli* that does not appear to perturb signaling in P. aeruginosa, suggesting a possible unappreciated level of complexity in PilT-PilJ signaling. Our data are consistent with a model in which PilT senses a surface through tension on the pilus fiber and relays this signal to modulate cAMP production.

## Results

### PilU levels significantly affect cAMP levels during surface attachment

To quantify cAMP levels during surface adaptation, the previously reported *PaQa* cAMP-responsive transcriptional reporter (4) was integrated onto the chromosome of *P. aeruginosa* PA14. This reporter is composed two fluorescent proteins, mKate2 and EYFP, under the control of two different promoters, *P_rpoD_* and *P_PaQa_*, respectively. *P_PaQa_* has been shown to be regulated by Vfr in a cAMP dependent manner and an increase in *P_PaQa_-eyfp* expression is correlated with an increase in cAMP. *P_rpoD_-mKate2* is used to normalize the EYFP levels for microscopy and used to gate on cells containing the reporter for flow cytometry (4). The *PaQa* reporter was integrated onto the chromosome at a neutral site of the *P. aeruginosa* PA14 chromosome using the mini-CTX1 system (4, 36). We validated this *PaQa* reporter using a mutant that is defective in cAMP production (Δ*cyaAB*) and a mutant lacking the phosphodiesterase that degrades cAMP (Δ*cpdA*), which results in over-production of cAMP (Figure S1A,B). After gating on single cells with *P_rpoD_*-mKate signal (gating cells with a ECD-A value of 1000 RFU or greater), the mean EYFP intensity was recorded and normalized to the WT signal. As expected, the Δ*cyaAB* mutant showed reduced levels of the cAMP reporter compared to the WT, while the Δ*cpdA* mutant showed an increased signal (Figure S1C).

While the PilT and PilU proteins both power retraction of T4P through ATP hydrolysis, these ATPases individually have unique roles in T4P dynamics and surface sensing. PilU is the accessory retraction motor for T4P in *P. aeruginosa*, whose function is dependent on the presence of PilT (37, 38). Both PilT and PilU hydrolyze ATP to power retraction but only PilT can interact with the platform protein PilC to coordinate PilA disassembly from extended T4P (32, 38). While PilT alone is able to retract pili bound to phage (as judged by phage sensitivity assays), PilU is required for T4P retraction that can pull the cell body along a surface to power twitching motility (TM) (14, 29, 30).

To better understand the respective contributions of PilT and PilU to surface-mediated cAMP production, we used single and double mutants and a series of phenotypic assays. The absence of PilT phenocopies a Δ*pilTΔpilU* double mutant in terms of TM, phage susceptibility and cAMP response (Figures 1A,B; and as reported previously by our group (29)). In contrast, a Δ*pilU* strain retains phage susceptibility due to the presence of PilT but shows an increase in surface-dependent cAMP response (Figure 1A,B; and as reported (15, 29, 33, 37–40)). Given that PilU is the only T4P protein whose deletion increases the level of surface-dependent cAMP and that this motor can only exert effects through PilT, we reasoned that PilT may be directly sensing the surface and relaying this signal to the Pil-Chp system, a model we probe in more detail below.

**Figure 1.**
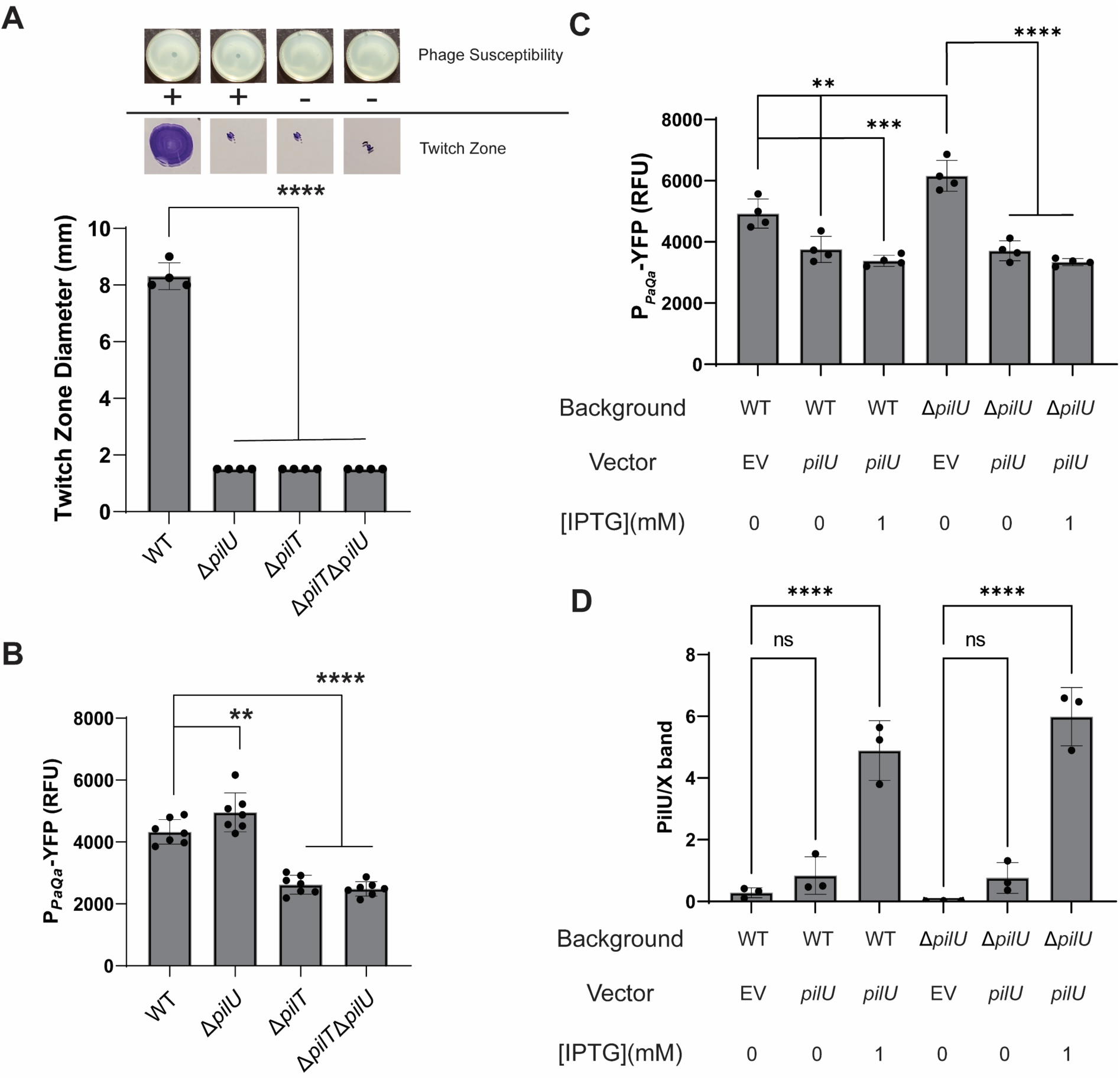
PilU levels affect surface-dependent cAMP production and T4P-related phenotypes. **A.** Images of phage sensitivity plates (top panel) and images of twitch zones stained with crystal violet (bottom panel). “+” indicates a phage susceptible strain and “–“ indicates a phage resistant strain. Below is the quantification of the twitch zone diameter for each strain. Data are from four biological replicates. **B.** Quantification of *PaQa* reporter as measured by flow cytometry after 5 hours of growth on agar. Data are from six biological replicates. **C.** Quantification of the *PaQa* reporter as measured by flow cytometry after 5 hours of growth for the WT and Δ*pilU* mutant expressing the *pilU* gene from a multicopy plasmid or carrying the empty vector (EV) control. Growth was on M8 agar supplemented with 1mM or no IPTG and the appropriate antibiotics. **D.** Quantification of the normalized PilU protein levels of the cells in panel C. Values were normalized to a cross-reacting band. Data are from three biological replicates. Bars and error bars in all panels are the mean and standard deviation and statistical significance was determined by one-way ANOVA followed by Tukey’s post hoc test, **P≤0.001, ***P≤0.0001, ****P≤0.00001; ns, not significant.

As mentioned above, PilT appears capable of retracting T4P under low loads like that of a phage bound to the pilus, but is unable to power twitching motility, which requires ATP hydrolysis from both PilT and PilU (37). If PilT attempts to retract a T4P filament bound to a surface without PilU present, we hypothesize that the motor would stall and enter a force induced conformational change, potentially due to improper ATP binding to, ATP hydrolysis of and/or ADP release from the hexamer (32). This PilT signaling model makes several predictions. First, a Δ*pilU* strain should show increased cAMP but only on a surface, a finding we have reported previously (15, 29, 39) and shown here (Figure 1B). Also, overexpression of PilU (a condition that is the opposite of deleting the *pilU* gene) should suppress the cAMP response.

To test this second prediction, we created the *pilU* expression plasmid, pVLT31-P_TAC_-*pilU,* and transformed this construct into the WT and Δ*pilU* mutant backgrounds with the *PaQa* reporter on the chromosome. cAMP was measured via flow cytometry with *pilU* expressed from a multi-copy plasmid for surface-grown bacteria (Figure 1C). Western blots were performed confirming excess PilU accumulation in all the analyzed strains (Figure 1D). All strains harboring the *pilU* overexpression construct had cAMP levels significantly lower than those with the vector control even in the absence of inducer. In the WT background, excess PilU significantly reduced the amount of cAMP when grown on a surface and the addition of inducer modestly further reduced the level of cAMP. The trends observed in the WT background were also observed in the Δ*pilU* mutant background. Interestingly, the level of cAMP production in the WT and Δ*pilU* backgrounds with the PilU construct were both significantly reduced from WT but not significantly different from each other. This observed decrease in surface-dependent cAMP is consistent with a model of PilT acting as a signaling protein during surface contact.

To determine whether the increase in cAMP production in the Δ*pilU* background was due to improper ATP hydrolysis during retraction, we expressed PilT-K136A (this mutation is in the Walker A box, WA, of ATPase domain) from a multicopy plasmid in the WT background. PilT-K136A is unable to bind ATP and we hypothesized that the incorporation of this monomer into the functional hexamer would lead to hexameric conformations similar to those that occur for the WT PilT during pilus retraction in absence of PilU. We observed a non-significant increase in cAMP in strains containing the PilT-K136A expression vector relative the empty vector control for the WT background (Figure S2, first 3 bars). This negative result is difficult to interpret because we do not know if the mutant protein is indeed incorporated into the motor.

### Mutations in PilT change the dynamics of surface-dependent cAMP induction

We previously showed that mutating critical residues in the Walker A and Walker B boxes of PilT, which abolish ATP binding and hydrolysis, respectively, also disrupt the surface-dependent cAMP response (29, 41). To investigate the role of PilT in surface sensing, these mutations, as well as other previously published mutations in the PilT protein of *P. aeruginosa* that affect the protein’s hexameric structure or retraction dynamics (32, 33, 42, 43), were inserted into the genome of *P. aeruginosa* PA14 at the gene’s native locus with the *PaQa* cAMP reporter on the chromosome at the neutral *attB* site. A list of mutations tested here with their characteristics, either previously published or determined in this report, are summarized in Table 1 and mapped onto the PilT protein structure (Figure 2A).

**Figure 2.**
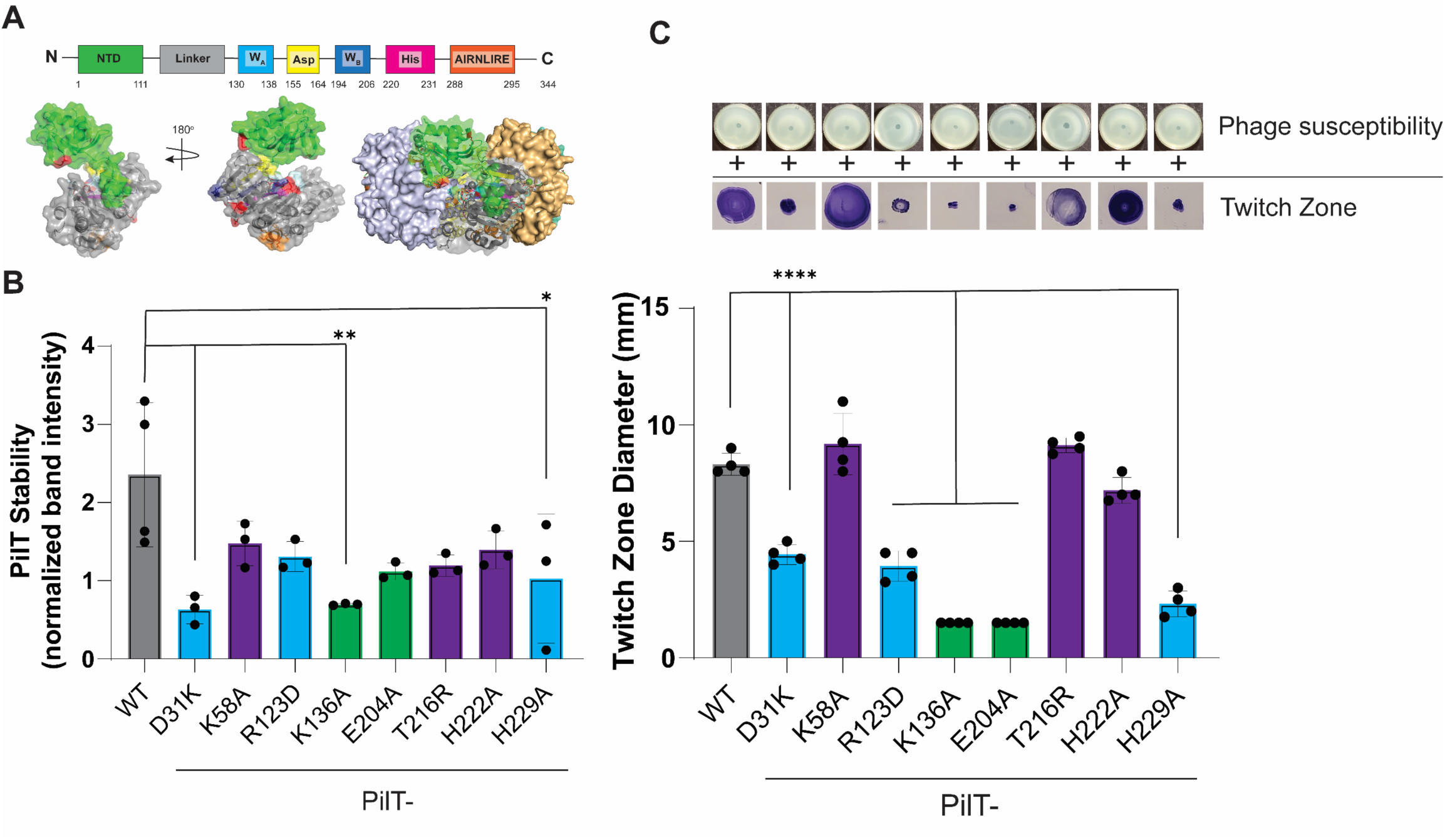
Characterization of PilT motor mutants. **A.** A schematic showing the domain architecture of PilT. Numbers represent the residue number of the beginning and ending of each domain. Below is the 3D structure of the *P. aeruginosa* PilT monomer and hexamer (PDB: 3jVV). **B.** Quantification of PilT protein levels via Western Blot analysis. The PilT band intensity from whole cells were normalized to a cross reacting band. Bars and errors bars represent the mean and SEM of 3 biological replicates. Data were analyzed by one-way ANOVA followed by Tukey’s post-test comparison. *, P<0.05, **, P<0.01, ***, P<0.001, ****, P<0.0001. Here and in panel C, strains that were able to twitch >75% of the WT have purple bars, strains that twitch between 25-75% of WT have blue bars, strains that twitch <25% WT have green bars. **C.** Assays for T4P function of *pilT* mutant strains. Images of phage sensitivity plates (top panel) and images of twitch zones stained with crystal violet (bottom panel). “+” denotes a phage susceptible strain and “-” denotes a phage resistant strain. The graph shows the quantification of twitch zone diameters for each strain. Bars and errors bars represent the mean and standard deviation of 4 biological replicates. Data were analyzed by one-way ANOVA followed by Tukey’s post-test comparison. ****, P<0.0001.

**Table 1.**
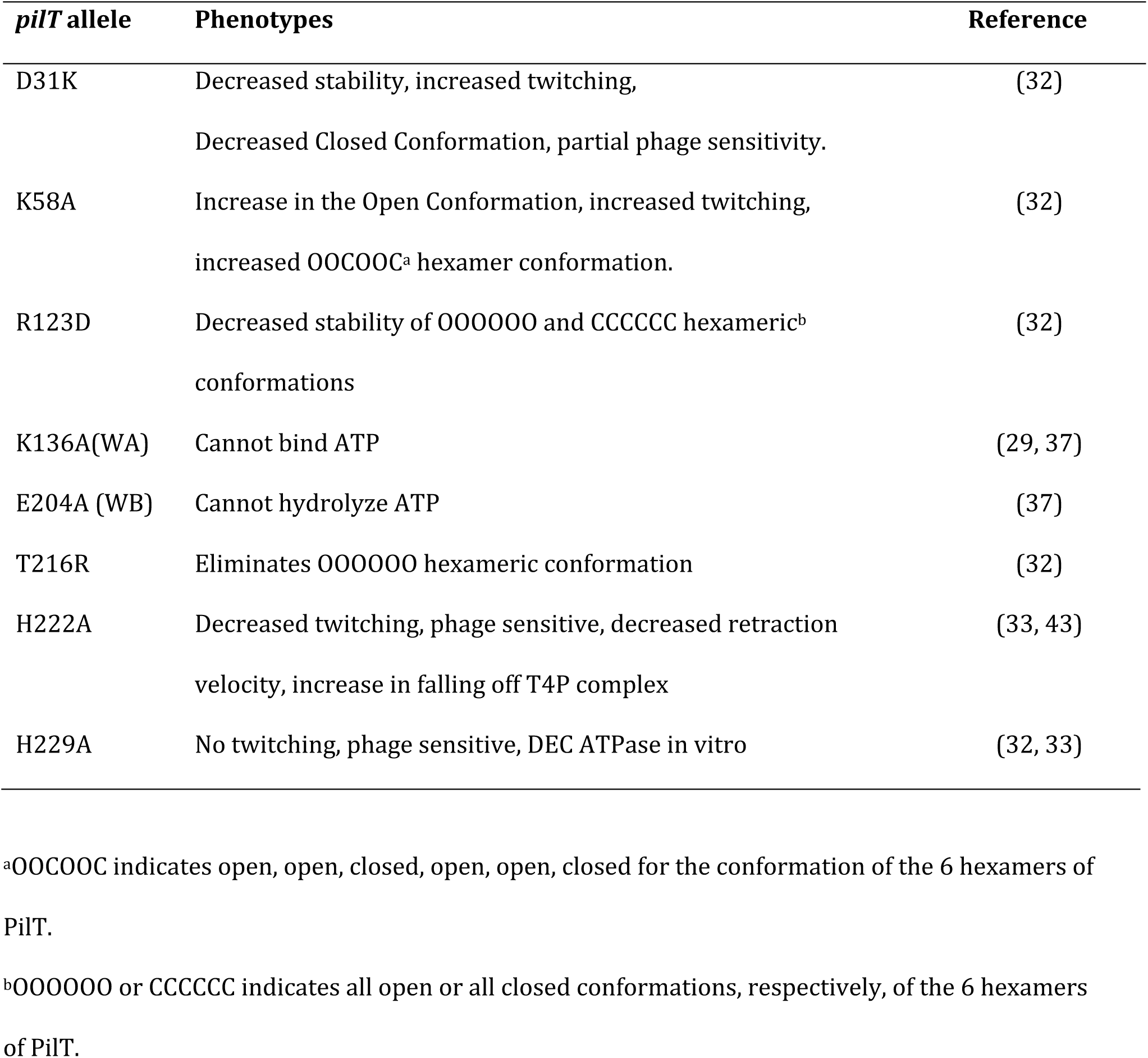
Mutant alleles characterized in this report.

For these mutants, we assessed protein stability, twitching motility, phage susceptibility and levels of cAMP when grown on a surface. The stability of each allele was assessed via Western blot and quantified by densitometry (Figure 2B). As previously reported, alleles E204A and D31K showed decreased stability (or perhaps antibody binding) (32). We also observed a significant decrease in the level of the protein with the PilT-H229A variant. The other alleles result in PilT levels that were reduced, but not significantly, relative to the level of the WT. The discrepancy in protein stability for some alleles could be due to the fact that previous reports used inducible promoters to express *pilT* on multi-copy plasmids and/or perhaps due to the fact the experiments were performed in the PAO1 strain (32, 33, 43). Here we produce PilT from a single copy with the gene’s native promoter to preserve endogenous regulation during surface sensing, and furthermore, to not perturb levels of PilU as the *pilU* gene is located directly downstream of the *pilT* gene.

To characterize the effects of these mutations on pilus retraction we performed twitching motility (TM) and phage susceptibility assays (Figure 2C). Twitching motility requires a fully functional PilT and PilU, while phage susceptibility requires only PilT (29, 44, 45). Despite the decrease in the level of the PilT proteins as measured by Western blot, all of the indicated PilT variants phenocopied previously published results in terms of twitching motility and phage susceptibility. Mutations that prevent ATP binding or hydrolysis (K136A, E204A; indicated in green) are unable to power TM but remain phage susceptible due to the presence of a functional PilU (29, 37). Mutating the first histidine in the His-box reduced TM while mutating the second histidine completely abolished TM (H222A, H229A; indicated in magenta and blue, respectively), however both alleles maintained phage susceptibility (32, 33). Mutations in the N-terminus of PilT and mutations that affect the overall hexameric structure of the PilT protein (D31K, K58A, R123D, T216R; blue and magenta bars) maintained phage susceptibility but the D31K and R123D mutations exhibited reduced TM (32).

Next, to capture the full dynamics of cAMP signaling during surface attachment, the first 6 hours of biofilm formation on the bottom of a glass bottom well was imaged using fluorescence microscopy. The average normalized fluorescent intensity per cell was plotted over time for cells harboring the *PaQa* reporter on the chromosome (2)(Figure 3A, Figure S3). All backgrounds initially start at the same level of intracellular cAMP (i.e., the lower levels associated with planktonic cells) and then begin to differ significantly for measured cAMP level within the first two hours. Throughout the time course, the Δ*pilT* (Figure 3A) and Walker box mutants (WA, PilT-K136A and WB, PilT-E204A; Figure 3A, Figure S3) maintained the lowest levels of cAMP. A strain carrying the PilT*-*D31K mutation (Figure S3) peaked early and then decreased after three hours of growth. The H229A allele maintained an intermediate level of cAMP relative to the other mutants. The remaining PilT alleles converged with the WT allele around hour 3 and maintained this trajectory until the end of the experiment although their cAMP levels differed from WT at most time points (Figure 3A, Figure S3).

**Figure 3.**
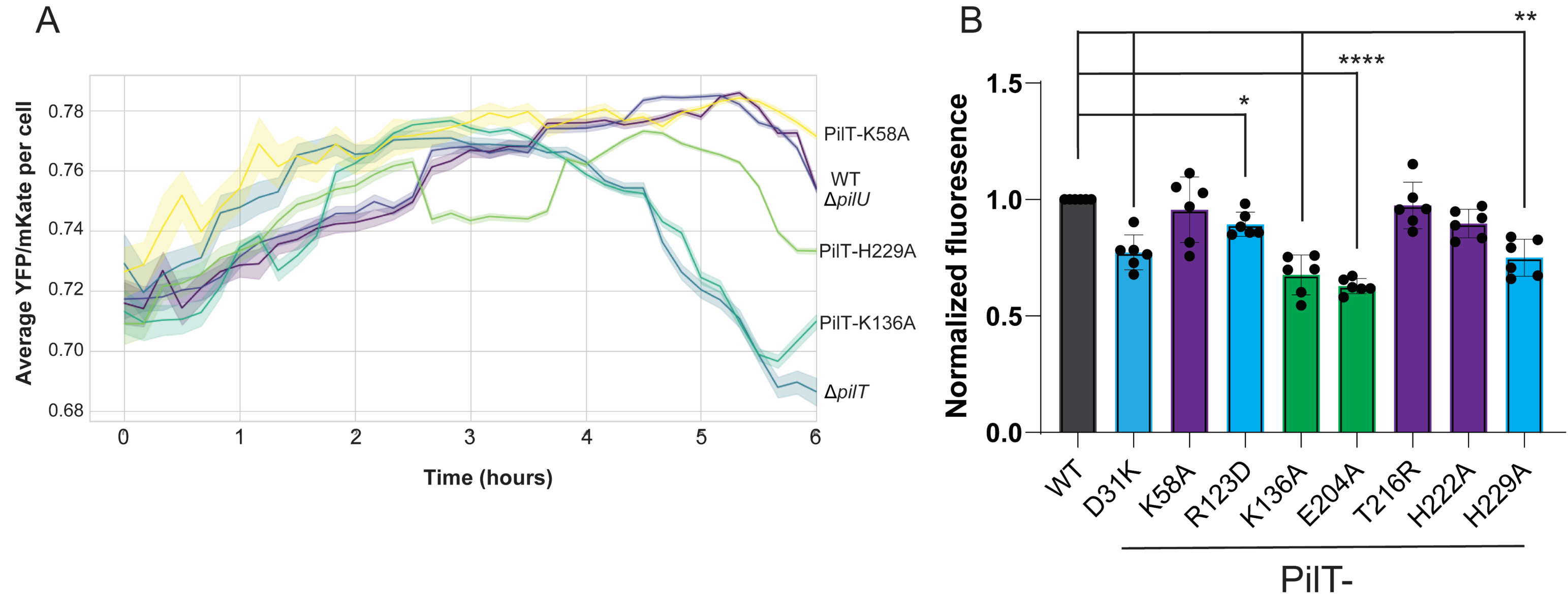
Measuring cAMP levels in strains carrying mutations in *pilT*. **A.** Graph depicting the average P*_PaQa_YFP*/P*_rpoD_mKate* per cell of selected strains during the first 6 hours of surface attachment in glass well dishes, as described in the text and Materials and Methods. Solid lines represent the mean YFP/mKate per cell and the shaded region represents the 95% confidence interval. At least 3 biological replicates were performed for each strain. A corresponding plot for all the *pilT* alleles described in this manuscript can be found in Fig S3. **B.** Cells were grown on an agar surface for 5 hours as described in the Materials and Methods and then analyzed by flow cytometer to quantify the amount of intracellular cAMP via the *PaQa* reporter. These values were then normalized by the WT value for that biological replicate. Bars and errors bars represent the mean and standard deviation of 6 biological replicates. Data were analyzed by one-way ANOVA followed by Tukey’s post-test comparison. **, P≤0.01, ***, P≤0.001, ****, P≤0.0001. Strains that were able to twitch >75% of the WT have purple bars, strains that twitch between 25-75% of WT have blue bars, strains that twitch <25% WT have green bars.

We noted that every genetic background that was unable to perform TM showed reduced levels of cAMP relative to WT (Figure 3B), with the exception of the Δ*pilU* mutant as shown above; this mutant background showed an increase in cAMP (Figure 1B) and as previously reported (15, 29, 39). cAMP was lowest in backgrounds that lacked the *pilT* gene followed by strains that expressed the Walker A (WA, PilT-K136A) and Walker B (WB, PilT-E204A) alleles of PilT. The PilT-D31K mutation resulted in a decrease in twitch zone and slight decrease in cAMP levels compared to that of WT. In contrast, PilT*-*H229A and R213D showed a significant decrease in twitching and cAMP levels. PilT-K58A, H222A, and T216R mutants displayed TM and cAMP not significantly different from WT.

As a control we measured cAMP levels in liquid grown cultures in the absence of a surface, all strains carrying these *pilT* alleles were not significantly different from the WT. As an additional control we grew the Δ*cpdA* mutant planktonically as well and it displayed high levels of cAMP (Figure S4A and (15)). We also showed that the cAMP measured in selected PilT alleles was PilJ-dependent, supporting the known role of PilJ in cAMP signaling (Figure S4B).

Together, these data show that mutations in various domains of the PilT motor alter cAMP signaling of bacteria grown on a surface, and these *pilT* alleles also impact pilus function by eliminating TM while retaining phage sensitivity, or alternatively, eliminating both TM and sensitivity to phage.

### Does PilT need to adopt both an ATP-(closed) and ADP-(open) bound state to support surface-dependent signaling?

Our previous work demonstrated that only the retractive force necessary for phage infection is necessary but not sufficient for surface-dependent cAMP induction (29). This conclusion was reached based on the observation that a Δ*pilU* mutant is phage susceptible, TM negative and has a cAMP level above that of the WT strain, while strains expressing the PilT-K136A (Walker A, WA) and PilT-E204A (Walker B, WB) mutations in a background with functional PilU is phage susceptible, TM negative but do *not* induce the cAMP response when grown on a surface (29).

While this observation could be due to a nuanced difference in the force threshold for cAMP induction versus phage infection, the lack of cAMP signaling in these PilT variants could also be due to the fact that these mutations limit the conformations that the PilT motor can adopt as a hexamer. That is, a fully functional PilT hexamer exists as a mixture of ATP-and ADP-bound states, and we hypothesized that a mixture of ATP-(closed) and ADP-bound (open) states of PilT might be necessary for cAMP signaling.

Given that the Walker A mutation prevents ATP-binding and the Walker B mutation prevents hydrolysis of ATP to ADP (32, 41), based on previous studies (32), locking the hexamer in either a fully unbound or fully ATP bound state, respectively, might interfere with surface signaling. Furthermore, given the ADP-bound state is structurally similar to nucleotide free state we reasoned that we may be able to observe the cAMP response if we expressed both Walker A and B mutants of PilT within the same cell.

To perform this experiment, we transformed a multicopy plasmid expressing the PilT*-*WA mutation (P_BAD_-*pilT*-K136A) in a background expressing the PilT*-*WB allele (PilT-E204A) integrated at its native locus with the *PaQa* reporter on the chromosome to measure the cAMP response. As shown in Figure S2 (last 3 bars), these mutations had no impact on cAMP levels, indicating that locking the PilT motor in these particular mixed conformations does not alter cAMP signaling.

### The retraction motor PilT binds to PilJ of the Pil-Chp system

The data presented so far are consistent with a model wherein PilT is required for the surface-dependent cAMP response. To determine how PilT might be influencing cAMP production, we screened for interactions between PilT and members of the Pil-Chp system using the Bacterial Adenylate Cyclase Two Hybrid System (B2H) and found an interaction between PilT and the protein at the top the Pil-Chp signaling system, PilJ (34) (Figure 4A,B). As a control we assessed binding between PilJ and the other retraction motor PilU and the extension motor PilB, but did not observe any such interaction (Figure 4A,B). We also assessed the interactions between PilT and the other components of the Pil-Chp system as well as other proteins that are known to influence surface attachment through T4P (Figure S5 and Table 2). We only detected robust interaction between PilT and PilJ, as well as the previously reported interaction between PilU and PilT (40).

**Figure 4.**
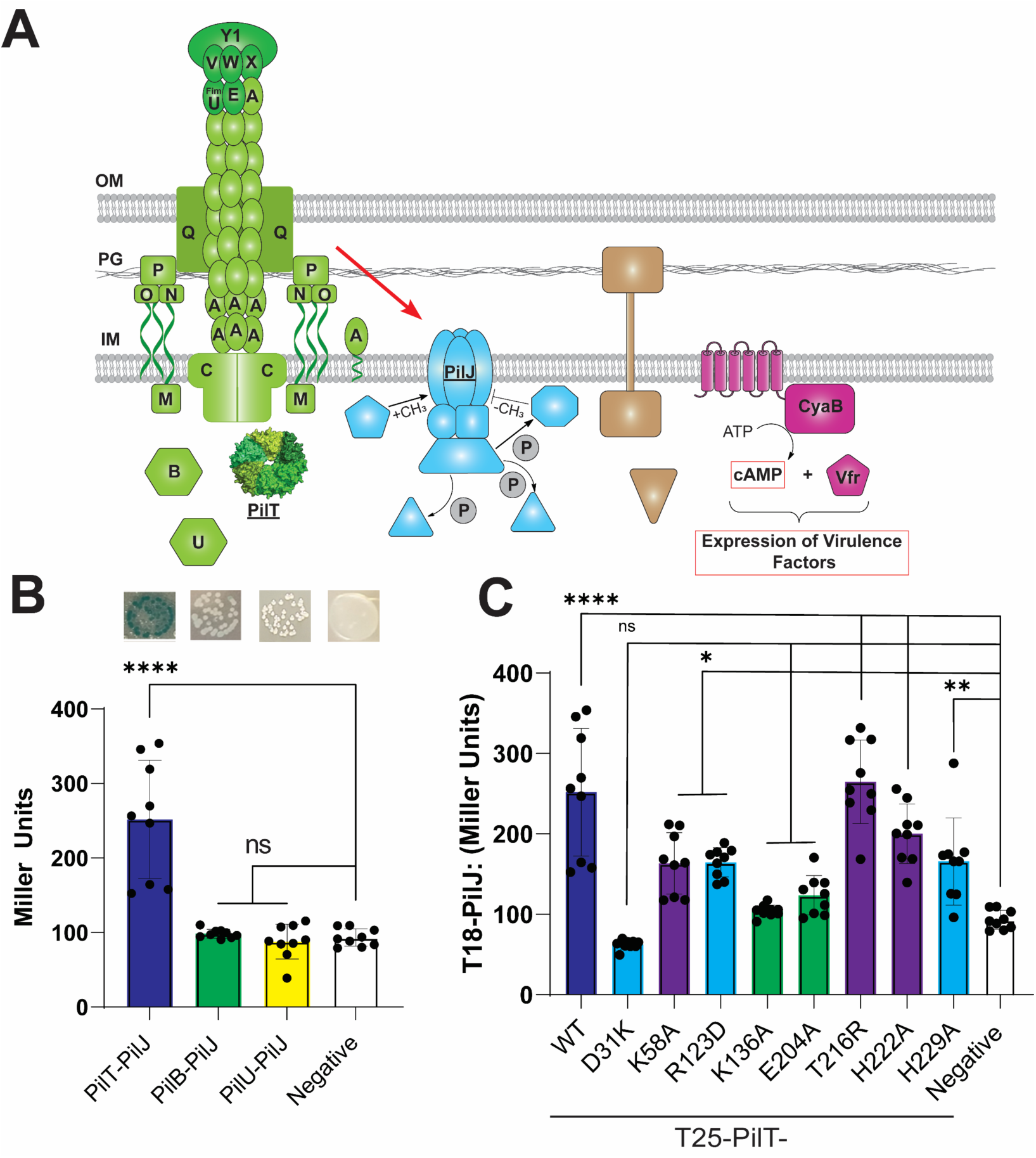
PilT interacts with PilJ. **A.** Schematic depicting the components of the TFP and cAMP-signaling pathway. **B.** Quantification of the B2H interaction between T4P motor proteins and PilJ in Miller Units. Shown are images of B2H colonies plated on X-gal plates (top of panel) and the interaction quantified (bottom of panel). Bars and errors bars represent the mean and standard deviation of 3 biological replicates. Data were analyzed by one-way ANOVA followed by Tukey’s post-test comparison. ns, not significant, ****, P<0.00001. **C.** Quantification of the level of interaction between different PilT mutants and PilJ using the B2H system. Bars and errors bars represent the mean and standard deviation of 3 biological replicates. Data were analyzed by one-way ANOVA followed by Tukey’s post-test comparison. ****, P<0.00001.

**Table 2.**
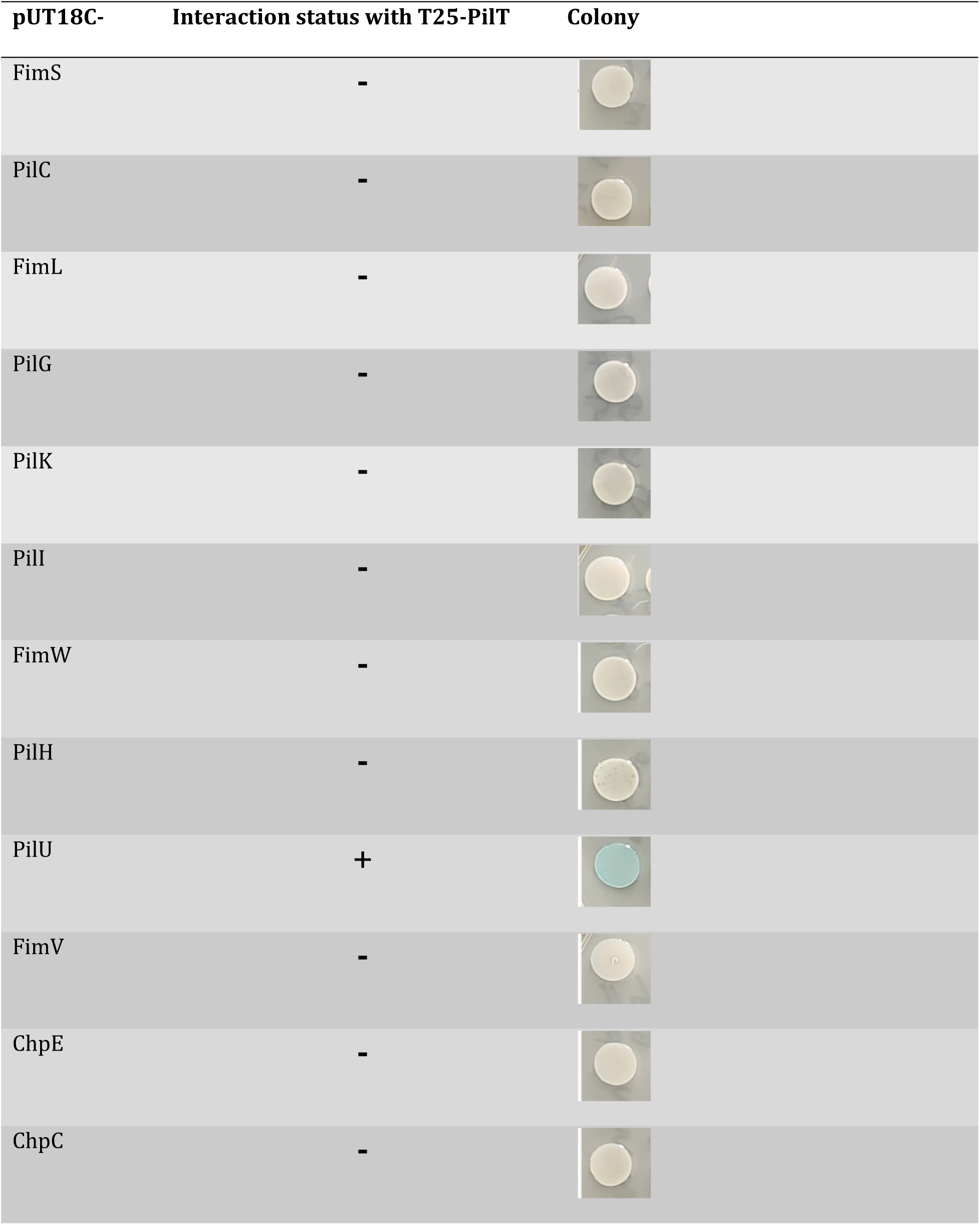
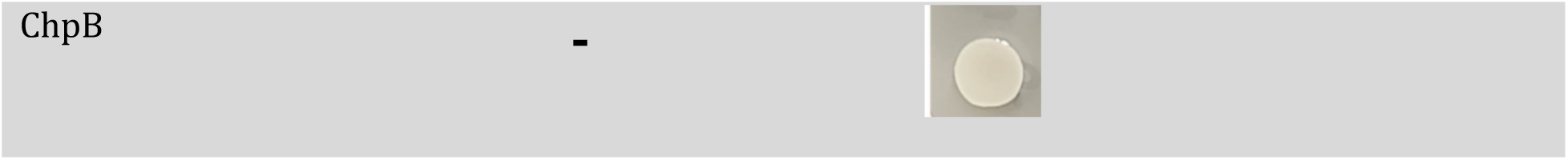
Interaction status with PilT using B2H assay.

To characterize this interaction with the *pilT* alleles described above, these mutants were cloned into the B2H system and the level of interaction with PilJ was measured via β-galactosidase activity. In general, PilT mutants that were able to perform TM and induce a surface-dependent cAMP response had higher levels of interaction with PilJ (Figure 4C).

We explore the consequences of the changes in interaction below.

### Associations between PilT-related phenotypes and cAMP signaling

We next examined associations between cAMP signaling and other measured phenotypes. Using our various *P. aeruginosa* mutant strains, we first plotted the diameter of the twitch zone for each allele versus the level of cAMP for surface-grown cells as measured by flow cytometry. Analyzing these data with a linear model we observed a highly significant, positive correlation between the twitch zone diameter and level of cAMP (Figure 5A). While this link between the production of cAMP and twitching motility is well known, the direct relationship between the levels of cAMP and the extent of TM, to our knowledge, has not been previously reported.

**Figure 5.**
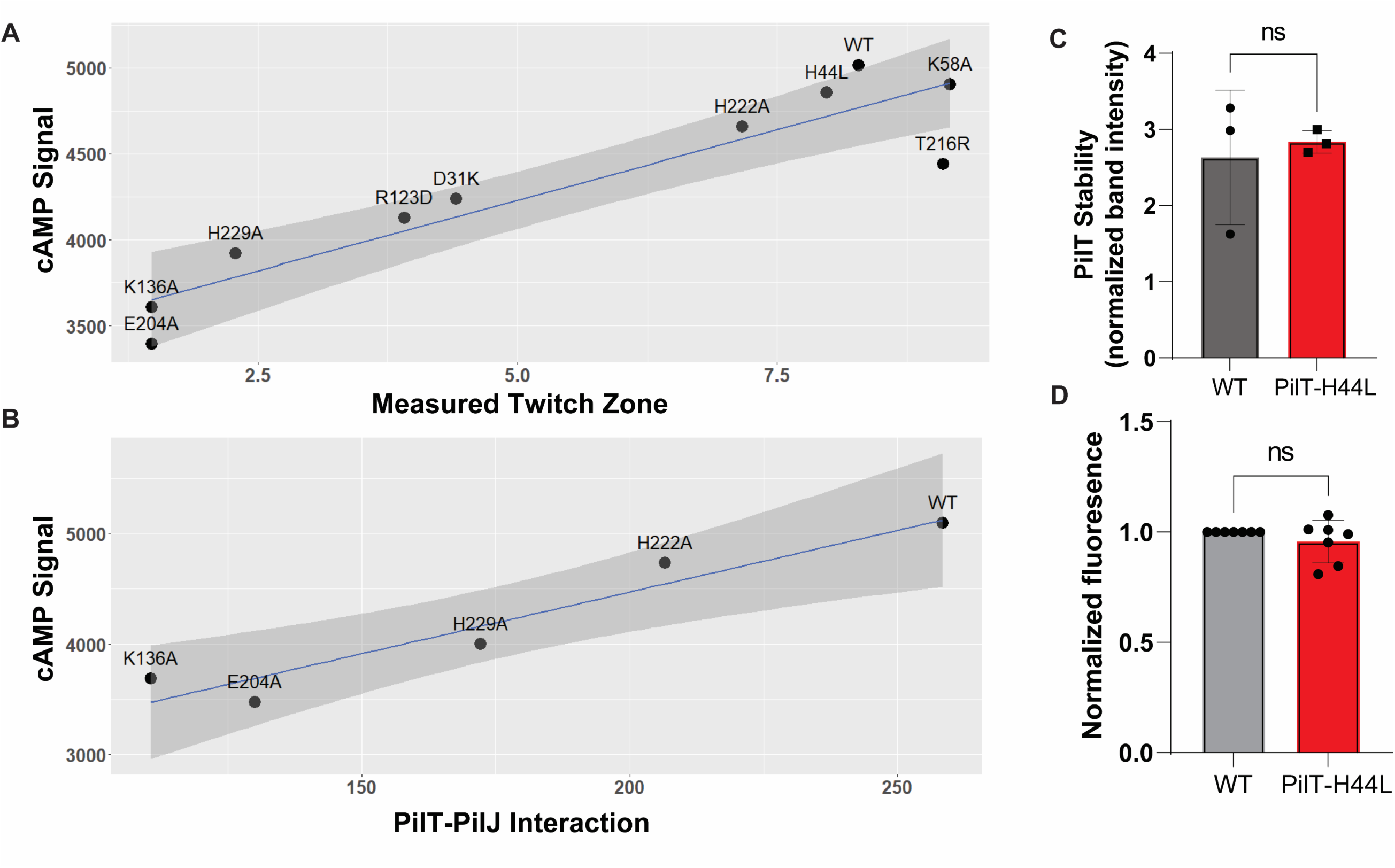
cAMP levels are positively associated with TM and the extent of PilT-PilJ interaction. **A.** A linear model depicting the relationship between twitching motility and surface induced cAMP production. Model is built using data from Figures 2, 3 and 4 using all tested alleles of *pilT* (R_2_=0.8565, Adjusted R_2_=0.8386, p-value=0.0001232). **B.** A linear model depicting the relationship between level of PilT-PilJ interaction as measured by the B2H system and surface induced cAMP production. Model is built using data from strains PilT-K136A, PilT-204A, PilT-H222A, PilT-H229A, and the WT strain (R_2_=0.9176, Adjusted R_2_=0.8902, pvalue=0.01029). **C.** Quantification of PilT level for the PilT-H44L mutant compared to the WT strain. PilT bands were normalized by a cross reacting band. Bars and errors bars represent the mean and standard deviation of 3 biological replicates compared to WT. Data are from 3 biological replicates and were analyzed by one-way ANOVA followed by Tukey’s post-test comparison. ns, not significant. **D**. Normalized fluorescence for WT and PilT-H44L strains. Values were normalized by the WT for each biological replicate. Bars and errors bars represent the mean and standard deviation of 7 biological replicates compared to WT. Data were analyzed by one-way ANOVA followed by Tukey’s post-test comparison. ns, not significant.

Importantly, using the data in Figures 3 and 4, we also observed a positive, significant relationship between cAMP levels and level of interaction between PilJ and PilT mutants that are completely or partially defective in ATPase activity (Figure 5B).

As a control, we also quantified the strength of interaction between the PilT mutants shown in Figure 5 and PilU using the B2H assay (Figure S6A). These values were then used with cAMP data to build a linear model to evaluate the relationship between PilT-PilU interaction strength and cAMP production (Figure S6B). We did not find any significant relationship between PilT-PilU interaction strength and cAMP levels when analyzing either all the PilT alleles or the mutations impacting ATPase activity, indicating that the relationship between PilT-ATPase variants and interaction strength with cAMP levels is specific to the PilT-PilJ interaction.

Together, these data indicate that the state of ATPase domain of PilT alters the interaction of this motor protein with the PilJ. Furthermore, this association between interaction strength of these ATPase mutants with PilJ and cAMP signaling suggests a mechanism whereby PilT ATPase activity could be linked to PilJ-mediated cAMP signaling, a possibility we discuss further below.

### Isolation of a mutation that disrupts PilJ-PilT interaction in E. coli using a B2H-based screen

To attempt to further understand the interaction between PilT and PilJ, and to assess the impact of this interaction on the influence of the cAMP response in *P. aeruginosa*, we screened for mutants of PilT that were able to perform twitching motility but no longer able to interact with PilJ using a B2H-based assay. A schematic describing the screening process can be found in Figure S7. Briefly, random mutations were introduced into the *pilT* sequence using error-prone PCR and then the mutant library cloned into the B2H backbone. This pool was then co-transformed with the WT PilJ construct and we picked white colonies indicating a loss of interaction with PilJ. The PilJ non-interacting alleles were then pooled and screened for the ability to retract pili by cloning this population of mutant *pilT* alleles into an expression vector and transforming this pool of alleles into the Δ*pilT* background of *P. aeruginosa*. The transformants were then screened for twitching motility. The *pilT* alleles that were able to twitch and did not interact with PilJ using a plate-based B2H assay were sequenced.

The extent of interaction of PilJ with candidate PilT alleles was then quantified using the B2H assay (Figure S8A), the stability of the proteins assessed by Western Blot (Figure S8B), and the mutations were mapped onto the PilT hexameric structure (Figure S8C). The majority of these mutations mapped to a patch on the surface of the N-terminal domain of the PilT protein.

As mentioned above, these mutant variants of PilT were checked for their stability via Western blotting, and unfortunately, the only stable allele was PilT-H44L (Figure S8B). The strain carrying the H44L showed levels of TM similar to the WT and the strain expressing this allele is phage susceptible (Fig S8D). The H44 residue maps to the N-terminal domain of PilT and should not impact ATPase activity. Although this allele does not interact with PilJ by B2H in *E. coli*, it appears to have phenotypes identical to the WT *P. aeruginosa* for twitching and phage susceptibility, and it produces cAMP levels that are not significantly different from the WT. These data suggest that the relationship between PilT and PilJ interaction and cAMP signaling may be complex, or that for this allele, the lack of interaction is *E. coli*-specific, points we discuss below.

## Discussion

Here, we examine the contribution of the T4P retraction motor PilT to surface sensing and present a model whereby PilT transmits a surface signal to the Pil-Chp system to activate cAMP production upon surface contact. Data presented here and from previous studies indicate that PilT is involved in sensing a surface, promoting at least minimal T4P retraction activity, and that PilT’s role may extended beyond its function in TFP retraction. We show that mutations in PilT that affect structure or T4P-related phenotypes also affect the surface-dependent cAMP response as measured through kinetic and endpoint assays. We also describe a novel interaction between PilT and PilJ of the Pil-Chp system. We then quantified the level of interaction between these PilT variants and PilJ using a B2H assay. A linear model showed a positive, significant correlation between cAMP level and twitching motility, and importantly, cAMP level and the strength or PilJ-PilT interaction for variants of PilT with known deficiencies in ATPase activity. We interpret this latter finding to mean that ATP binding and hydrolysis is critical, not only for retraction activity, but for PilT to bind and transmit a signal to PilJ. Furthermore, the interaction strength between the PilT ATPase mutants and PilJ is correlated with an increase in cAMP. We did not observe this correlation when examining alleles of PilT with full ATPase activity or the interaction between PilT and PilU. Together, these data support a model whereby the retraction motor PilT senses the surface during T4P retraction and relays this signal to PilJ.

We and many others have shown that loss of PilU function results in an increase in surface-dependent cAMP levels (15, 29, 39). We propose that when the PilU accessory retraction motor is absent during retraction, PilT enters a force induced conformational change, perhaps through stalling of the motor while attempting to retract a bound pilus, in turn transmitting a surface signal to the Pil-Chp system. Our model makes several predictions that have been confirmed in previous publications (15, 29, 39) and this study. First, we demonstrated that cAMP levels when grown on a surface are dependent on the presence of PilT and PilJ. Second, the absence of PilU leads to elevated cAMP levels, a finding made by others and confirmed here (15, 29, 39). Lastly, our data show that excess PilU, has the opposite effect, reducing the surface-dependent cAMP response.

We also observed a strong, positive correlation between twitching ability as measured through twitch zone diameter and cAMP production for multiple mutants when analyzed with a linear model. This finding is consistent with previous studies that demonstrated that twitching motility requires cAMP production, and that at the single cell level, oscillations of T4P activity and cAMP are highly correlated (39). While cAMP production is dependent on T4P activity, this second messenger activates a positive feedback loop whereby Vfr and cAMP positively regulate Fims-AlgR, which in turn leads to increased expression of minor pilins and the number of active T4P complexes per cell. This positive regulatory loop is part of the rapid surface adaptation response of *P. aeruginosa*, and we now know that part of this signaling cascade is initiated by the retraction motor PilT. Thus, PilT appears to play integrated roles in surface sensing and the control of TM in response to surface inputs.

To complement our candidate mutant approach for studying the PilT-PilJ interaction, we performed a genetic screen to identify PilT variants that could promote TM despite a defect in interaction with PilJ. This screen yielded two interesting findings. First, most of the mutations mapped to a region near the N-terminus of PilT with a known role in hexamer formation. The N-terminus of PilT binds to the C-terminus of the adjacent monomer to form the hexamer. This interface also undergoes the major conformational change during ATP binding and also makes contacts that facilitate the O and C conformations. Thus, these mutations could impact PilT hexamer protein assembly and/or function. Unfortunately, most of these PilT variants were unstable thus we could not unwind whether the lack of signaling was due to reduced levels of the PilT protein, or the inability of these mutant proteins to interact with PilJ.

We did isolate one allele of *pilT* that was able to perform TM but did not interact with PilJ in *E. coli* using a B2H assay; for this allele (H44L) we still observed WT surface-dependent cAMP production in *P. aeruginosa*. This mutation was near the N-terminal region of PilT and not in proximity to any parts of the protein thought to contribute to its ATPase activity. Consistent with this idea, this allele was able to perform TM at a level similar to that of WT, indicating that this mutant variant does not have a defect in its ATPase activity. A similar phenotype was observed for the PilT-D31K allele, which retains ATPase activity (32). This PilT-D31K allele, which also maps to the N-terminus of PilT, was able to perform TM at WT levels and induce cAMP production when on a surface, but had very low levels of interaction with PilJ when measured through the B2H assay. Thus, it is possible that mutations at the N-terminus of PilT can impact its ability to interact with PilJ in the B2H, but perhaps not impact PilT-PilJ interaction when this allele is expressed in *P. aeruginosa* in the presence of the rest of the T4P machinery or in the context of the hexamer. We believe it is important to acknowledge that while our findings here allow us to posit a model connecting T4P to surface sensing and cAMP signaling, we still lack key pieces of information to build a model which explains all of the current data. We look forward to interrogating our model further.

We believe our findings are consistent with previous studies, as T4P motors as signaling proteins is not unprecedented. For example, *Mxyococcus xanthus* requires T4P for exopolysaccharide (EPS) production as well as a type of surface-based motility known as S-motility. Researchers performed a suppressor screen for EPS production in a T4P-deficient background. Mutations in the T4P assembly ATPase PilB were isolated that led to the production of EPS without S-motility. A Walker-A mutation in PilB phenocopied this mutation and was dominant over the WT allele (46). Consistent with our studies of T4P/PilY1 regulating surface-dependent cdG signaling (3), these data link a T4P and motor function to second messenger signaling, and may represent a more general strategy whereby T4P (and perhaps other pili families) serve double duty as adhesins and signal transduction machinery.

Others have suggested that the surface signal is transmitted from the T4P pilin, PilA, to PilJ to activate cAMP production. Although we have previously shown no correlation between PilA-PilJ binding strength and cAMP production (29) this does not exclude a role for PilA signaling through a different mechanism. Regardless of the extent of signaling through PilA, the pilin remains a critical part of our motor signaling model, as the motor relies on the pressence of a pilus fiber to extend and bind to the surface to create tension during PilT retraction. Overall, we have presented evidence for a new model of surface signal sensing and transduction through the T4P motor PilT and we will continue to investigate the mechanism by which this occurs.

## Materials and Methods

### Strains and media

*Pseudomonas aeruginosa* UCBPP PA14 was used as the WT strain. Mutations were made in this background using *E. coli* S17-1 λpir. *E. coli* BTH101 was used for Bacterial Adenylate Cyclase Two Hybrid assays. Strains used in this study are listed in Supplemental Table S1. Bacterial strains were routinely cultured in 5ml of lysogeny broth (LB) medium or plated on 1.5% agar with antibiotics when necessary. Tetracycline (tet) was used at 15ug/ml for *E. coli* and 120 ug/ml during *P. aeruginosa* selection and maintained with 75ug/ml. Gentamicin (Gm) was used at 30 µg/ml for *P. aeruginosa* and 10 µg/ml for *E. coli*. Carbenicillin (Cb) was used at 250 ug/ml for *P. aeruginosa* and 100 ug/ml for *E. coli*. Kanamycin (Kan) was used at 50 ug/ml for *E. coli*. M8 minimal salts medium supplemented with MgSO4 (1mM), glucose (0.2%) and casamino acids (0.5%) was used for all assays. Plasmids were induced with either 0.2% arabinose for P_BAD_ promoter induction or 0.5 mM isopropyl-D-thiogalactopyranoside (IPTG) was added to agar or liquid media for P_TAC_ promoter induction unless otherwise stated. β-galactosidase activity from B2H assays was visualized using plates supplemented with 5-bromo-4-chloro-3-indolyl-β-D-galactopyranoside (X-Gal; 40 μg/ml).

### Construction of mutant strains and plasmids

Plasmids used in this study are listed in Supplemental Table S2 and primers are listed in Supplemental Table S3. Plasmids were constructed using Gibson assembly of purified PCR products. Chromosomal mutations were made using homologous recombination with the pMQ30 vector. Insertions at neutral sites in the *P. aeruginosa* genome were made using the mini-Tn7 vector (47, 48) and the mini-CTX1 vector (36). Resistance markers were removed using the pFLP2 plasmid followed by sucrose counter selection (36). Point mutations were generated using QuikChange ® site-directed mutagenesis followed by Gibson assembly. Expression vectors were generated using Gibson assembly of purified PCR products into pMQ72 or pVLT31 and then transformed into *P. aeruginosa* or *E. coli* using electroporation.

### Twitching motility

Twitching motility plates were made using M8 medium supplemented with casamino acids, MgSO4, glucose and 1% agar. Plates were inoculated from liquid cultures using a sterile toothpick plunged through the agar to the bottom of the plate. Plates were incubated for 24 hours at 37C and then 24 hours at room temperature. The agar was then removed and the twitch zones were stained with 0.1% crystal violet. Images were obtained and the twitch zone diameter was measured twice using a ruler.

### Phage plaque assay

Phage susceptibility assays were performed in (60 x 15 mm) plates using M8 medium supplemented with casamino acids, MgSO4, glucose, and 1% agar. 1ml of 0.5% M8 molten agar was then inoculated with 50 ul from a *P. aeruginosa* overnight culture. This mixture was poured over the solidified 1% M8 agar to form a bacterial lawn. After solidifying, 2ul of phage DMS3_vir_ lysate was pipetted onto the bacterial lawn and incubated for 24 hours at 37C.

### Bacterial Adenylate Cyclase Two Hybrid assays

The B2H system from Euromedex (49) was used to assess protein-protein interactions in *E. coli* BTH101. Alleles of *pilT* and other T4P proteins were cloned into the pKT25 vector and PilJ along with other Pil-Chp proteins were cloned into the pUT18 and pUT18C vectors. A pair of pKT25 and pUT18/UT18C vectors were then co-transformed into *E. coli* BTH101. To visualize the interaction, transformants were plated on LB agar containing Cb, Kan, X-Gal (5-bromo-4-chloro-3-indolyl-β-D-galactopyranoside) (40 g/ml) and IPTG (isopropyl-D-thiogalactopyranoside) (0.5 mM) and incubated at 30C until an interaction was observed through the transformation of X-Gal to a blue pigment or until the negative control began to produce a blue pigment. To quantify the level of interaction between proteins, transformants were plated on LB agar with Cb, Kan and IPTG. After incubation at 30C, cells were harvested and β-galactosidase assays were performed as previously described (49).

### Protein detection and quantification

Strains were grown in M8 liquid medium supplemented with arabinose or IPTG and grown at 37C for 6 hours. Whole cell lysates were prepared as previously described (50). Cultures were OD normalized to 1 and an equal volume was resolved on either a 12% or 10% polyacrylamide gel. Proteins were then transferred to a nitrocellulose membrane and probed with either anti-PilT or anti-PilU antisera. Detection of proteins was performed using fluorescence detection with IR-Dye®-labeled fluorescent secondary antibodies and imaged using the Odyssey CLx Imager (LICOR Biosciences, Inc., Lincoln, NE). Quantification of protein bands was performed using Image Studio Lite software (LICOR Biosciences, Inc., Lincoln, NE). Protein levels were then normalized by a cross-reacting band.

### Flow cytometry measurements

Bacterial strains harboring the *PaQa* reporter on the chromosome were subcultured into liquid M8 medium supplemented with glucose, casamino acids, and MgSO4 and incubated at 37C until an OD of 0.5 was reached, ∼3 hours. Gm, Tet, IPTG, or arabinose was added to the liquid medium when indicated. 200ul of the culture was then spread onto M8 agar plates and allowed to incubate for 5 hours at 37C. Cells were then harvested from these plates, washed, diluted, and analyzed on a Beckman Coulter Cytoflex S. FlowJo software version 10.8.1 was used to gate on populations of single cells that had mKate fluorescence. The EYFP fluorescence from the *P_PaQa_* promoter was then measured on the gated population. A workflow of the gating strategy can be found in Figure S9.

### Microscopy experiments using 8-well dishes

Bacterial strains were subcultured in liquid M8 supplemented with glucose, MgSO4, and casamino acids after being cultured overnight in liquid LB at 37C. After reaching an OD600∼0.5, cultures were diluted 1:100 into fresh, liquid M8 and then 300ul was used to fill a glass bottom chamber (Cellvis 8 chambered cover glass system). The chamber was then mounted on a Nikon Ti Eclipse epifluorescence microscope in an environmental chamber set to 37C. Images of at least 3 FOV were taken every 5 minutes for the first 8 hours of surface attachment. Images were then analyzed using a python script, which can be found at https://github.com/GeiselBiofilm.

### Statistical analysis

Data visualization and statistical analysis were performed in GraphPad Prism 9 (version 9.2.0). Linear mixed models were built in R (v4.0.2) and visualized using ggplot2 (v3.3.2). The script used to perform the analysis can be found at https://github.com/GeiselBiofilm.

## Data Accessibility Statement

All code is available on Guthub at https://github.com/GeiselBiofilm.

## Acknowledgements

We thank Katie Forest and Lori Burrows for helpful discussions, Lori Burrows for providing antibodies, Tom Hampton for help with linear modelling, Zdenek Svindrych for help with microscopy, Ko-Wei Liu, Matthew James, Stacie Stutt, and Gary Ward for help with flow cytometry, and Sherry Kuchma, Shanice Webster, Berenike Maier, and Amruta Karbelkar for helpful discussions. This work was funded by the NIH (RO1 R01AI143730 to GAO) and BioMT through NIH NIGMS grant P20-GM113132.

