## Supplemental Figures for "Evidence for the Type IV Pili Retraction Motor PilT as a Component of the Surface Sensing System in *Pseudomonas aeruginosa*"

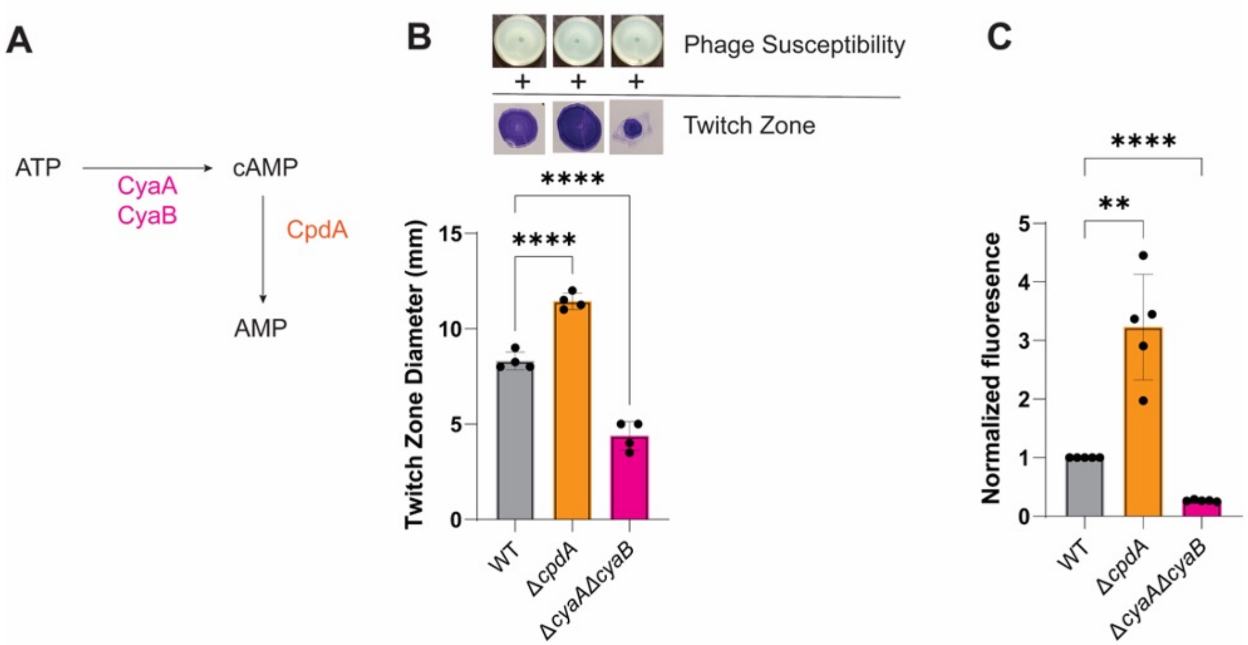


**Figure S1. Analyzing cAMP levels to validate the reporter. A.** A diagram representing the enzymes that make and degrade cAMP in *P. aeruginosa*. **B.** Phage susceptibility assay and twitching motility assay for the WT, Δ*cpdA,* and Δ*cyaAB* mutant backgrounds. Images of phage susceptibility plates (top panel, "+" indicates that strains are susceptible to phage infection) and twitching motility zones stained by crystal violet (middle panel) are above the quantification of the twitch zone diameter. Bars and errors bars represent the mean and standard deviation of 4 biological replicates when compared to the WT. Data were analyzed by one-way ANOVA followed by Tukey's post-test comparison. ****, P≤0.00001, **, P≤0.001. **C.** Normalized fluorescence for WT and the Δ*cpdA,* and Δ*cyaAB* mutant strains. Values were normalized to the WT for each biological replicate. Bars and errors bars represent the mean and standard deviation of 3 biological replicates compared to WT. Data were analyzed by one-way ANOVA followed by Tukey's post-test comparison. ****, P≤0.00001, **, P≤0.01.

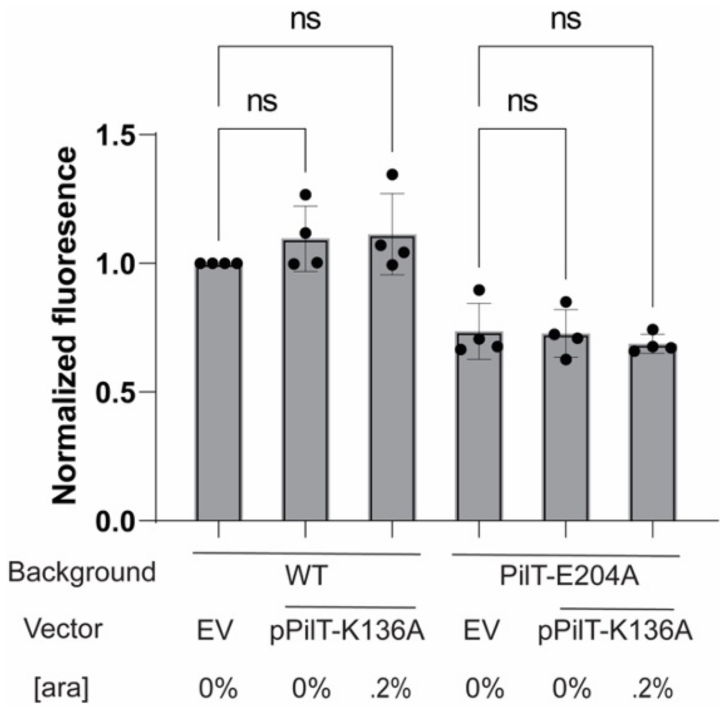


**Figure S2. Co-expressing the Walker A (WA)/Walker B (WB) mutations does not impact cAMP signaling.** Quantification of cAMP in WT (left) or PilT-E204A (right) strains carrying a plasmid expressing the PilT-K136A variant or the empty vector (EV) control, supplemented with 0 or 0.2% arabinose. Values were normalized to the WT for each corresponding biological replicate. Bars and errors bars represent the mean and standard deviation of 4 biological replicates. Data were analyzed by one-way ANOVA followed by Tukey's post-test comparison. ns, not significant.

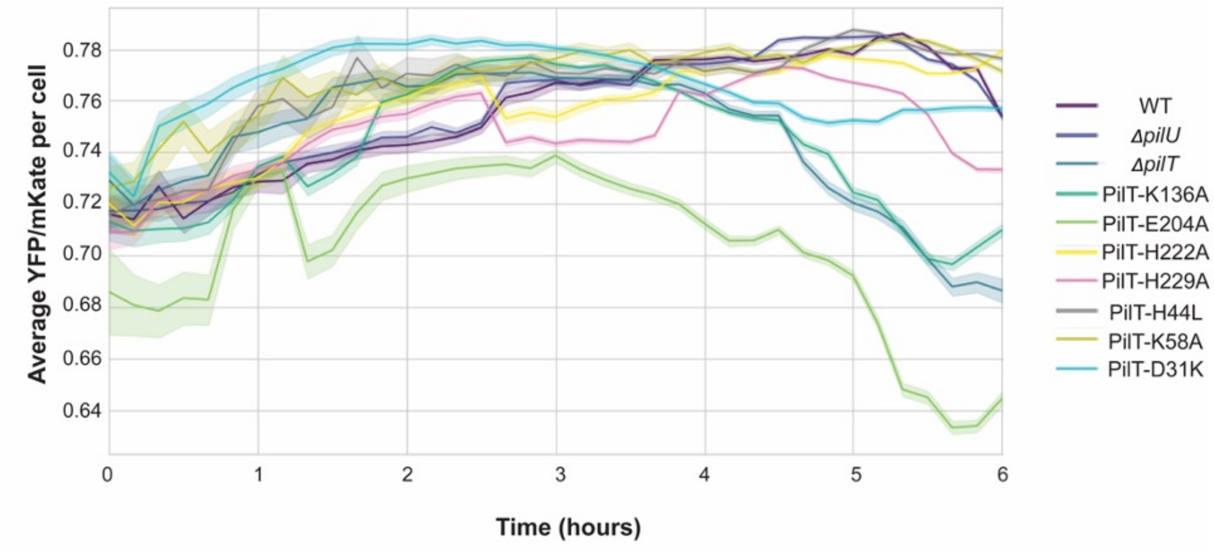


**Figure S3. Time course measuring cAMP levels.** Graph depicting the average $P_{PaQa}$*YFP*/$P_{rpoD}$*mKate* per cell of selected strains during the first 6 hours of surface attachment in glass well dishes, as described in the text and Materials and Methods. Solid lines represent the mean YFP/mKate per cell and the shaded region represents the 95% confidence interval. At least 3 biological replicates were performed for each strain.

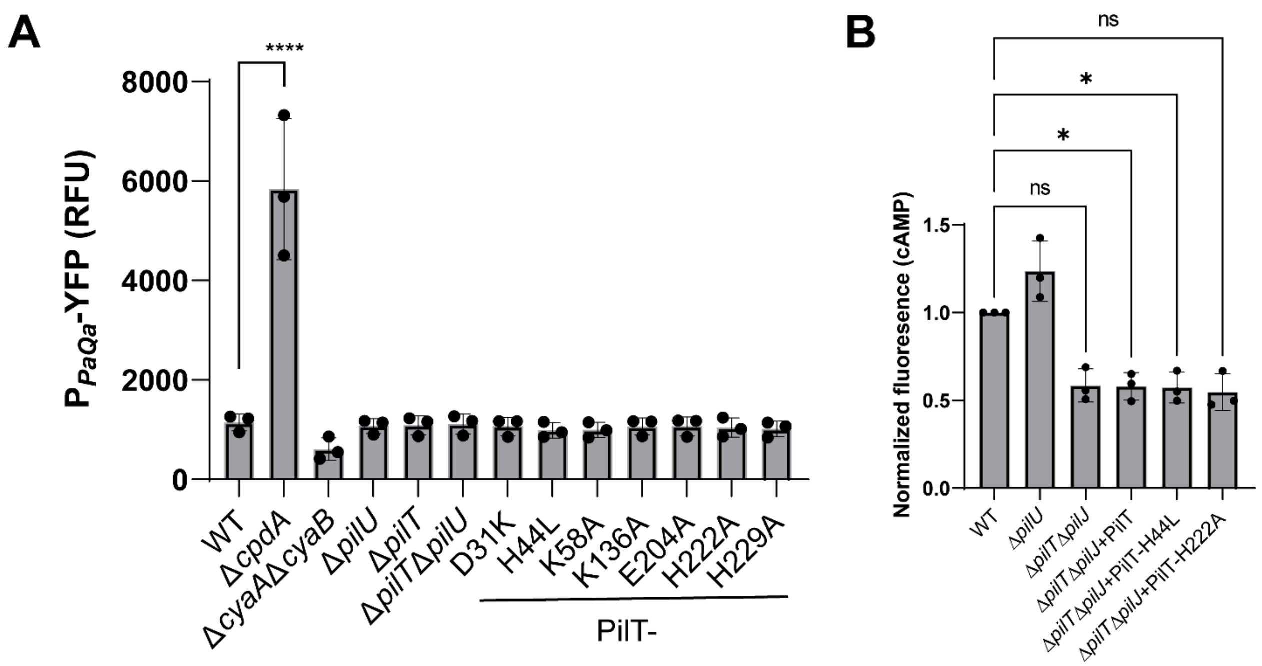


**Figure S4. cAMP signaling is dependent on a surface and PilJ. A.** Intracellular cAMP levels as measured by the *PaQa* reporter for the indicated mutants grown in planktonic culture. Bars and errors bars represent the mean and standard deviation of 3 biological replicates when compared to the WT. Data were analyzed by one-way ANOVA followed by Tukey's post-test comparison. ****, P≤0.00001, ns, not significant. **B**. Signaling via selected PilT alleles is PilJ dependent. Quantification of cAMP as measured by the PaQa reporter using flow cytometry. Data were analyzed by one-way ANOVA followed by Tukey's post-test comparison. *, P≤0.05, ns, not significant.

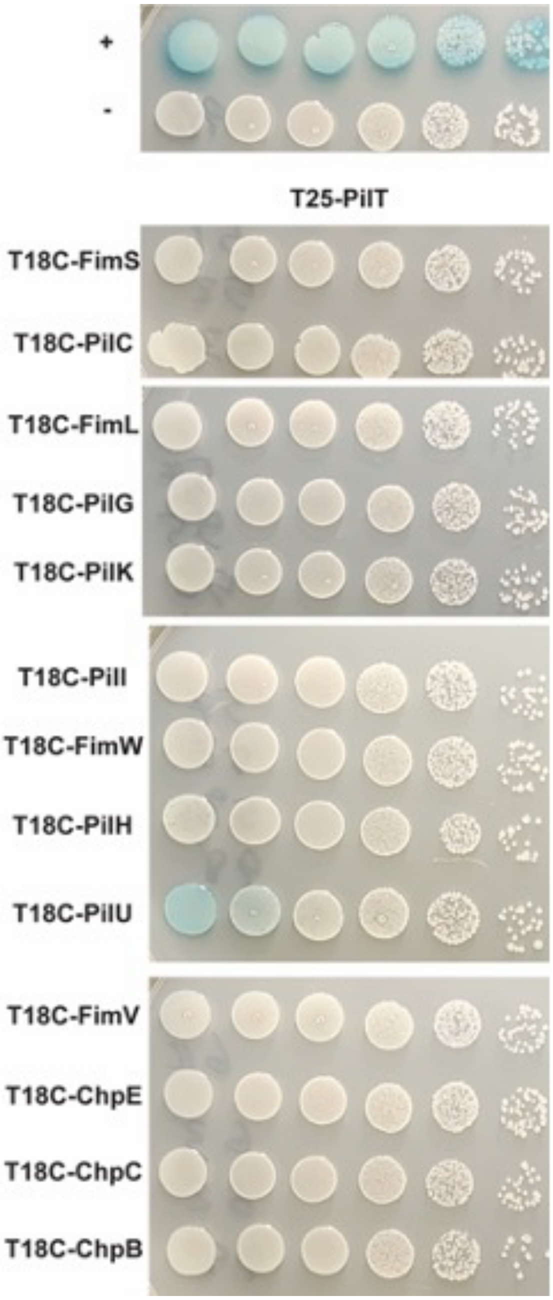


**Figure S5**. **PilT does not interact with other members of the Pil-Chp system as measured by B2H.** Images of plates from the B2H assay assessing interactions with PilT. As described in the Materials and Methods, BTH101 cells were co-transformed with the pUT25-PilT plasmid and the pUT18C plasmid fused to different components of the Pil-Chp system. Transformants were then serially diluted and plated on agar containing X-Gal and the appropriate antibiotics. The only observed interaction was between PilT and PilU. Positive (+) and negative (-) B2H controls are shown at the top of the figure.

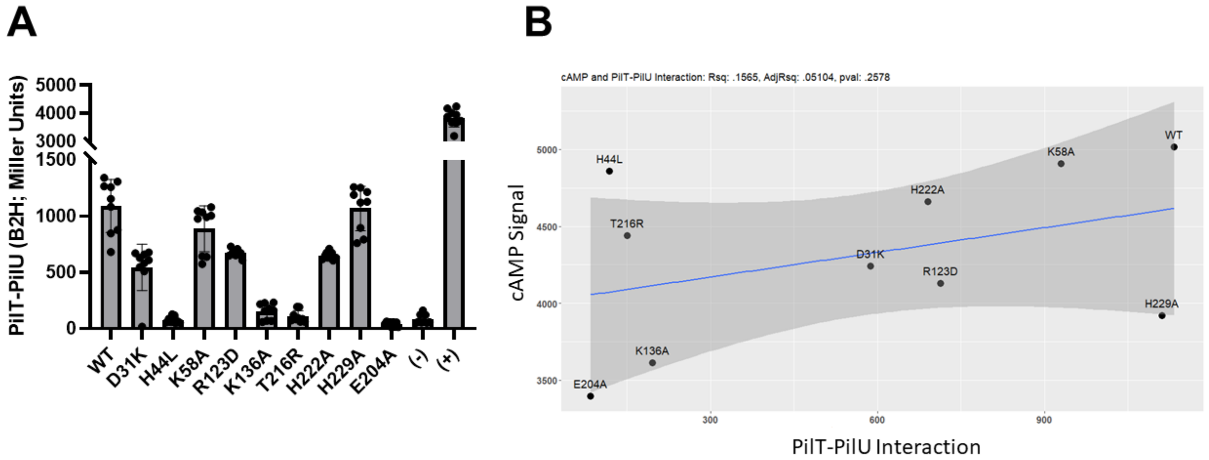


**Figure S6. PilU and PilT interaction strength and cAMP**. **A.** Quantification of the interaction between PilT mutants and PilU as measured by the B2H assay. Bars and errors bars represent the mean and standard deviation of 3 biological replicates when compared to the WT. **B.** A linear model depicting the relationship between level of PilT-PilU interaction as measured by the B2H system and surface induced cAMP production. The model was built using data from all mutants of PilT in R ($R^2$: 0.1565, Adjusted $R^2$: 0.05104, p-value: 0.2578). When only considering the ATPase mutants, a similar not significant result is obtained ($R^2$: 0.5406, Adjusted $R^2$: 0.3875, p-value: 0.1569).

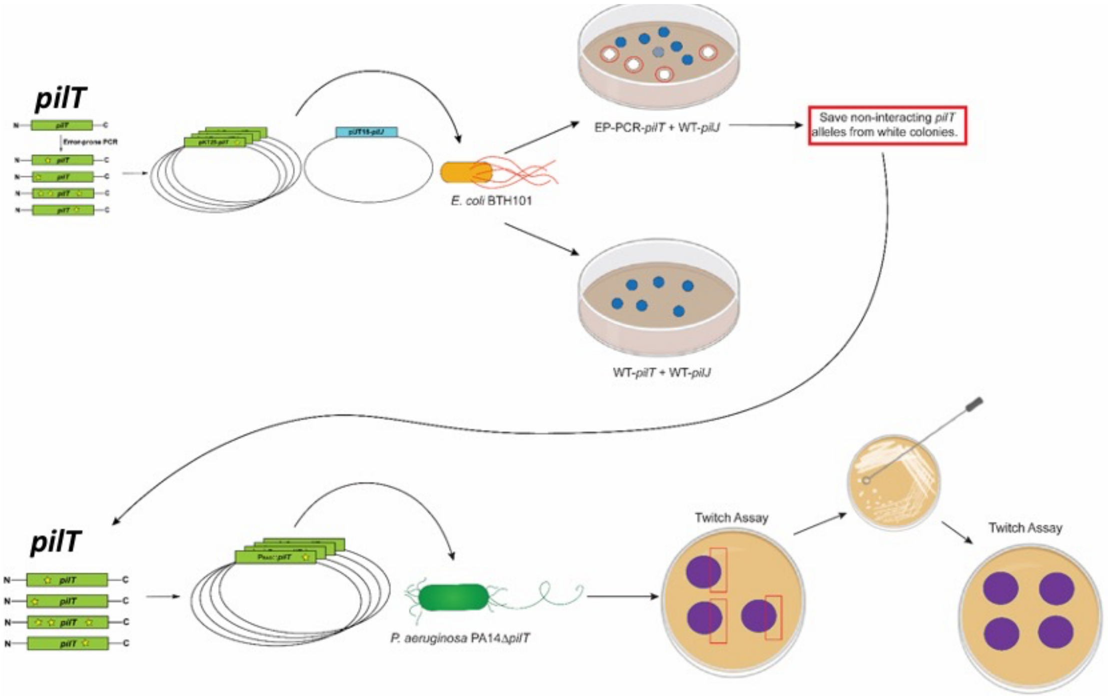


**Figure S7. Isolating a mutation that disrupts PilT-PilJ interaction.** A schematic depicting the screening process to isolate functional *pilT* alleles that no longer bind to PilJ in the B2H assay.

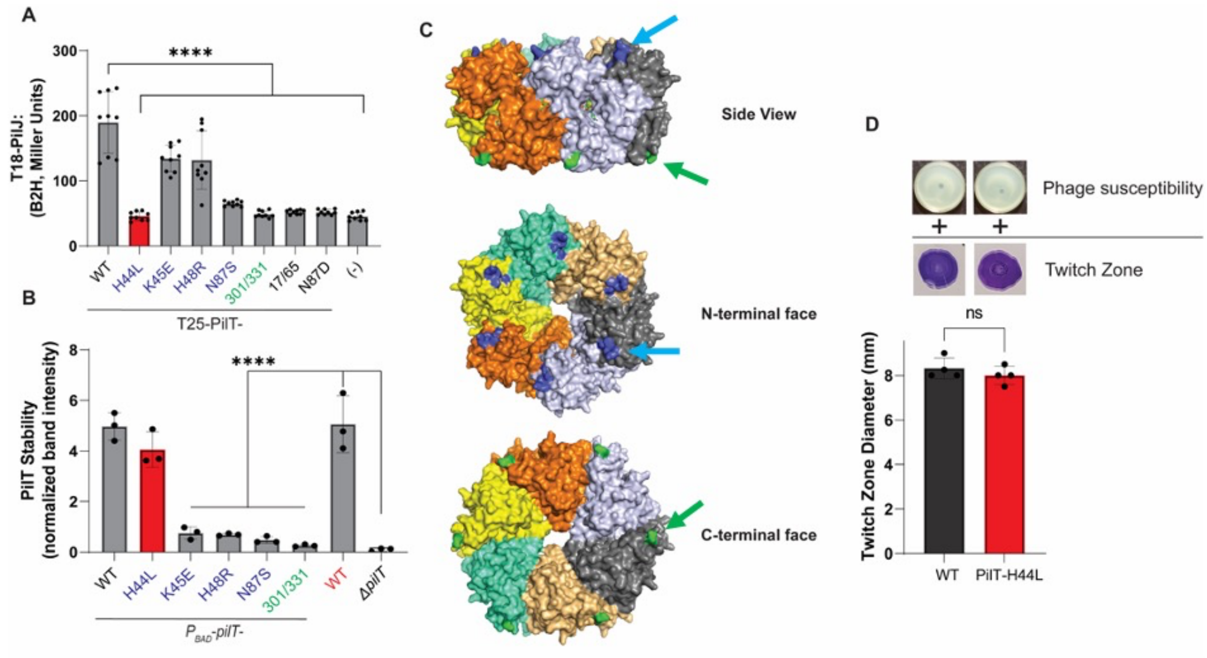


**Figure S8. Analysis of mutants from the genetic screen**. **A**. Quantification of the level of interaction between different PilT mutants and PilJ using the B2H system. Mutations in blue map to the blue patch on the N-terminal face of PilT shown in panel C. Mutations in green localize to the C-terminal face as shown in panel C. Bars and errors bars represent the mean and standard deviation of 3 biological replicates. The red bar represents the PilT-H44L mutant. Data were analyzed by one-way ANOVA followed by Tukey's post-test comparison. ****, $P \leq 0.00001$. **B.** PilT levels expressed from a multi-copy plasmid and quantified via Western blot. Bars and errors bars represent the mean and standard deviation of 3 biological replicates. All PilT intensity values were normalized by a cross-reacting band. Data were analyzed by one-way ANOVA followed by Tukey's post-test comparison. ****, $P < 0.00001$. **C**. Hexameric structure of PilT (PDB: 3jVV) side view (top panel), N-terminal face (middle panel), and C-terminal face (bottom panel) are imaged. Each monomer is in a different color. The mutations listed in blue of panels A and B cluster on the N-terminal face of the hexamer and highlighted in blue with a blue arrow pointing at it. The mutations listed in green form a patch on the C-terminal face of the hexamer and colored green, indicated by a green arrow. **D.** T4P assays to assess PilT-H44L function. Images of phage susceptibility plates (top panel) and twitching motility zones stained by crystal violet (middle panel) are above the quantification of the twitch zone diameter. Bars and errors bars represent the mean and standard deviation of 3 biological replicates when compared to the WT. Data were analyzed by one-way ANOVA followed by Tukey's post-test comparison. ns, not significant.

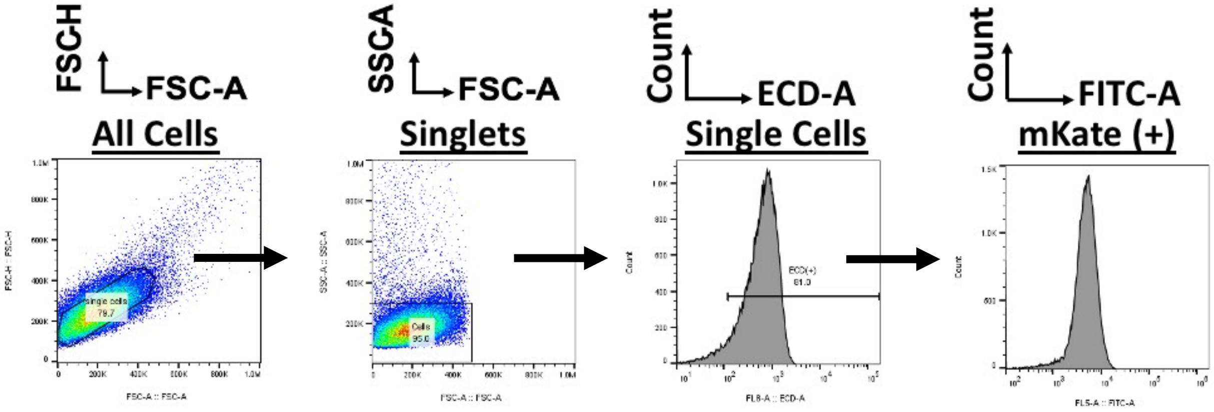


**Figure S9. Gating strategy for flow cytometry assays.** Single cells were first selected by gating on the FSC-H to FSC-A scatter plot and then gating by size in the SSC-A by FSC-A scatter plot. This population was then further gated to only include mKate positive cells by gating cells with a ECD-A value of 1000 RFU or greater. The resulting population was then used to calculate the geometric mean and standard deviation of the FITC-A population.
