## Supplemental Tables for "Evidence for the Type IV Pili Retraction Motor PilT as a Component of the Surface Sensing System in *Pseudomonas aeruginosa*"

### Supplementary Table S1. Strains used in this study.

| Strain name | Relevant genotype, description | Source |
|---|---|---|
| *E. coli* strains | | |
| DH5α | *supE44 ΔlacU169(f80lacZΔM15) hsdR17 thi-1 relA1 recA1* | Life Technologies |
| S17-1(λpir) | *thi pro hsdR- hsdM+ ΔrecA RP4-2::TcMu-Km::Tn7* | (1) |
| BTH101 | *F-, cya-99, araD139, galE15, galK16, rpsL1 (Str r), hsdR2, mcrA1, mcrB1* | (2) |
| SM10 | *thi thr leu tonA lacY supE recA::RP4-2-Tc::Mu Km λpir* | (1) |
| *P. aeruginosa* strains (SMC#) | | |
| 232 | PA14 wild type (WT) | (3) |
| 7302 | Δ*pilT;* unmarked in-frame deletion | (4) |
| 7304 | Δ*pilU;* unmarked in-frame deletion | (5) |
| 8856 | Δ*pilT*Δ*pilU;* unmarked in-frame deletion | (6) |
| 9652 | Δ*pilT*Δ*pilJ;* unmarked in-frame deletions | This study |
| 6707 | Δ*cyaA*Δ*cyaB;* unmarked in-frame deletions | (5) |
| 6851 | Δ*cpdA;* unmarked in-frame deletion | (5) |
| 9462 | PA14 *attTn7::P1-lacZ, attB::*$P_{PaQa}$*-eyfp,* $P_{rpoD}$*-mKate2;* Tetracycline-resistance ($Tc^r$) cassette | This study |
| 9463 | Δ*pilU attB::*$P_{PaQa}$*-eyfp,* $P_{rpoD}$*-mKate2;* $Tc^r$ | This study |
| 9464 | Δ*pilT attB::*$P_{PaQa}$*-eyfp,* $P_{rpoD}$*-mKate2;* $Tc^r$ | This study |

| | | |
|---|---|---|
| 9465 | Δ*pilT*Δ*pilU attB::*$P_{PaQa}$*-eyfp,* $P_{rpoD}$*-mKate2;* Tc$^{r}$ | This study |
| 9466 | Δ*cpdA attTn7::P1-lacZ, attB::*$P_{PaQa}$*-eyfp,* $P_{rpoD}$*-mKate2;* Tc$^{r}$ | This study |
| 9467 | Δ*cyaA*Δ*cyaB attTn7::P1-lacZ, attB::*$P_{PaQa}$*-eyfp,* $P_{rpoD}$*-mKate2;* Tc$^{r}$; Tc$^{r}$ | This study |
| 9653 | *pilT-D31K attTn7::P1-lacZ, attB::*$P_{PaQa}$*-eyfp,* $P_{rpoD}$*-mKate2;* Tc$^{r}$ | This study |
| 9654 | *pilT-H44L attTn7::P1-lacZ, attB::*$P_{PaQa}$*-eyfp,* $P_{rpoD}$*-mKate2;* Tc$^{r}$ | This study |
| 9655 | *pilT-K58A attTn7::P1-lacZ, attB::*$P_{PaQa}$*-eyfp,* $P_{rpoD}$*-mKate2;* Tc$^{r}$ | This study |
| 9656 | *pilT-R123D attTn7::P1-lacZ, attB::*$P_{PaQa}$*-eyfp,* $P_{rpoD}$*-mKate2;* Tc$^{r}$ | This study |
| 9665 | *pilT-K136A attTn7::P1-lacZ, attB::*$P_{PaQa}$*-eyfp,* $P_{rpoD}$*-mKate2;* Tc$^{r}$ | This study |
| 9657 | *pilT-E204A attTn7::P1-lacZ, attB::*$P_{PaQa}$*-eyfp,* $P_{rpoD}$*-mKate2;* Tc$^{r}$ | This study |
| 9658 | *pilT-T216R attTn7::P1-lacZ, attB::*$P_{PaQa}$*-eyfp,* $P_{rpoD}$*-mKate2;* Tc$^{r}$ | This study |
| 9659 | *pilT-H222A attTn7::P1-lacZ, attB::*$P_{PaQa}$*-eyfp,* $P_{rpoD}$*-mKate2;* Tc$^{r}$ | This study |
| 9660 | *pilT-H229A attTn7::P1-lacZ, attB::*$P_{PaQa}$*-eyfp,* $P_{rpoD}$*-mKate2;* Tc$^{r}$ | This study |
| 9661 | PA14 *attTn7::P1-lacZ, attB::*$P_{PaQa}$*-eyfp,* $P_{rpoD}$*-mKate2* | This study |
| 9662 | Δ*pilU attB::*$P_{PaQa}$*-eyfp,* $P_{rpoD}$*-mKate2* | This study |
| 9663 | Δ*pilT*Δ*pilJ attB::*$P_{PaQa}$*-eyfp,* $P_{rpoD}$*-mKate2,* Tc$^{r}$ | This study |
| 9664 | Δ*pilT*Δ*pilU attB::*$P_{PaQa}$*-eyfp,* $P_{rpoD}$*-mKate2* | This study |

**Supplementary Table S2. Plasmids used in this study.**

| Plasmid name | Description | Source |
|---|---|---|
| pMQ72 | Vector for cloning in yeast, *E. coli*, and *P. aeruginosa* with arabinose-inducible gene expression system; Gentamycin-resistance ($Gm^r$) cassette | (7) |
| pMQ30 | Shuttle vector for cloning in yeat and allelic exchange in Gram-negatives; $Gm^r$ | (7) |
| miniCTX1 | Suicide vector for integration into the *Pseudomonas attB* site on the chromosome; Tetracycline-resistance ($Tc^r$) cassette | (8) |
| pMQ56-mTn7 | Suicide vector for insertion of the mini-Tn7 element from pUT18-mTn7-Gm into the *attTn7* site on the chromosome of *P. aeruginosa* | (6) |
| pFLP2 | For removal of antibiotic resistance cassettes flanked by FRT sites with Flipase; sacB for plasmid counterselection; Carbenicillin-resistance ($Cb^r$) cassette | (9) |
| pUT18C | BACTH vector for fusions with the C-terminus of the T18 fragment of *cyaA*, $Cb^r$ | Euromedex |
| pUT18 | BACTH vector for fusions with the N-terminus of the T18 fragment of *cyaA*, $Cb^r$ | Euromedex |
| pKT25 | BACTH vector for fusions with the C-terminus of the T25 fragment of *cyaA*, Kanamycin resistance ($Kn^r$) cassette | Euromedex |
| pKNT25 | BACTH vector for fusions with the N-terminus of the T25 fragment of *cyaA*, Kanamycin resistance ($Kn^r$) cassette | Euromedex |

| | | |
|---|---|---|
| pKT25-zip | BACTH positive control, leucine zipper of GCN4 fused to T25, $Kn^r$ | Euromedex |
| pUT18C-zip | BACTH positive control, leucine zipper of GCN4 fused to T18, $Cb^r$ | Euromedex |
| pVLT31 | Vector for cloning in *E. coli* and *P. aeruginosa* with IPTG-inducible gene expression system; $Tc^r$ | (10) |
| pMQ72-*pilT* | For arabinose-inducible expression of *pilT* in *P. aeruginosa;* $Gm^r$ | This study |
| pMQ72-*pilT-H44L* | For arabinose-inducible expression of *pilT-H44L* in *P. aeruginosa;* $Gm^r$ | This study |
| pMQ72-*pilT-H222A* | For arabinose-inducible expression of *pilT-H222A* in *P. aeruginosa;* $Gm^r$ | This study |
| pVLT31-*pilU* | For IPTG-inducible expression of *pilU* in *P. aeruginosa;* $Tc^r$ | This study |
| pKT25-*pilT* | *pilT* gene cloned into pKT25; $Kn^r$ | This study |
| pKT25-*pilT-D31K* | *pilT-D31K* gene cloned into pKT25; $Kn^r$ | This study |
| pKT25-*pilT-K58A* | *pilT-K58A* gene cloned into pKT25; $Kn^r$ | This study |
| pKT25-*pilT-K136A* | *pilT-K136A* gene cloned into pKT25; $Kn^r$ | This study |
| pKT25-*pilT-E204A* | *pilT-E204A* gene cloned into pKT25; $Kn^r$ | This study |
| pKT25-*pilT-H222A* | *pilT-H222A* gene cloned into pKT25; $Kn^r$ | This study |
| pKT25-*pilT-H229A* | *pilT-H229A* gene cloned into pKT25; $Kn^r$ | This study |
| pKT25-*pilT-H44L* | *pilT-H44L* gene cloned into pKT25; $Kn^r$ | This study |
| pKT25-*pilT-K45E* | *pilT-K45E* gene cloned into pKT25; $Kn^r$ | This study |
| pKT25-*pilT-H48R* | *pilT-H48R* gene cloned into pKT25; $Kn^r$ | This study |
| pKT25-*pilT-N87S* | *pilT-N87S* gene cloned into pKT25; $Kn^r$ | This study |

| | | |
|---|---|---|
| pKT25-*pilT-N87D* | *pilT-N87D* gene cloned into pKT25; $Kn^r$ | This study |
| pKT25-*pilT-D17G/E65K* | *pilT-D17G/E65K* gene cloned into pKT25; $Kn^r$ | This study |
| pKT25-*pilT-M301T/R331H* | *pilT-M301T/R331H* gene cloned into pKT25; $Kn^r$ | This study |
| pUT18-*pilJ* | *pilJ* gene cloned into pUT18; $Cb^r$ | |
| pUT18C-*fimS* | *fimS* gene cloned into pUT18C; $Cb^r$ | This study |
| pUT18C-*pilC* | *pilC* gene cloned into pUT18C; $Cb^r$ | This study |
| pUT18C-*fimL* | *fimL* gene cloned into pUT18C; $Cb^r$ | This study |
| pUT18C-*pilG* | *pilG* gene cloned into pUT18C; $Cb^r$ | This study |
| pUT18C-*pilK* | *pilK* gene cloned into pUT18C; $Cb^r$ | This study |
| pUT18C-*pilI* | *pilI* gene cloned into pUT18C; $Cb^r$ | This study |
| pUT18C-*fimW* | *fimW* gene cloned into pUT18C; $Cb^r$ | This study |
| pUT18C-*pilH* | *pilH* gene cloned into pUT18C; $Cb^r$ | This study |
| pUT18C-*pilU* | *pilU* gene cloned into pUT18C; $Cb^r$ | This study |
| pUT18C-*fimV* | *fimV* gene cloned into pUT18C; $Cb^r$ | This study |
| pUT18C-*chpE* | *chpE* gene cloned into pUT18C; $Cb^r$ | This study |
| pUT18C-*chpC* | *chpC* gene cloned into pUT18C; $Cb^r$ | This study |
| pUT18C-*chpB* | *chpB* gene cloned into pUT18C; $Cb^r$ | This study |
| pMQ72-*pilT-H44L* | For arabinose-inducible expression of *pilT-H44L* in *P. aeruginosa;* $Gm^r$ | This study |
| pMQ72-*pilT-K45E* | For arabinose-inducible expression of *pilT-K45E* in *P. aeruginosa;* $Gm^r$ | This study |
| pMQ72-*pilT-H48R* | For arabinose-inducible expression of *pilT-H48R* in *P. aeruginosa;* $Gm^r$ | This study |

| | | |
|---|---|---|
| pMQ72-*pilT-N87S* | For arabinose-inducible expression of *pilT-N87S* in *P. aeruginosa;* $Gm^r$ | This study |
| pMQ72-*pilT-M301T/R331H* | For arabinose-inducible expression of *pilT-M301T/R331H* in *P. aeruginosa;* $Gm^r$ | This study |
| miniCTX1-*PaQa* | For integration of the *PaQa* reporter into the *attB* site of the chromosome of *P. aeruginosa*. $P_{PaQa}$-*eyfp*, $P_{rpoD}$-*mKate2* | This study |
| pMQ56-mTn7-*P1-lacZ* | For integration of the cAMP transcriptional reporter into the *attTn7* site on the chromosome of *P. aeruginosa*. *P1-lacZ* | (6) |
| pMQ30-*pilT-D31K* | For performing allelic exchange at the native locus of the *pilT* gene in *P. aeruginosa* PA14 to introduce the mutation *pilT-D31K* | This study |
| pMQ30-*pilT-H44L* | For performing allelic exchange at the native locus of the *pilT* gene in *P. aeruginosa* PA14 to introduce the mutation *pilT-H44L* | This study |
| pMQ30-*pilT-K58A* | For performing allelic exchange at the native locus of the *pilT* gene in *P. aeruginosa* PA14 to introduce the mutation *pilT-K58A* | This study |
| pMQ30-*pilT-R123D* | For performing allelic exchange at the native locus of the *pilT* gene in *P. aeruginosa* PA14 to introduce the mutation *pilT-R123D* | This study |
| pMQ30-*pilT-E204A* | For performing allelic exchange at the native locus of the *pilT* gene in *P. aeruginosa* PA14 to introduce the mutation *pilT-E204A* | This study |

| | | |
|---|---|---|
| pMQ30-*pilT-T216R* | For performing allelic exchange at the native locus of the *pilT* gene in *P. aeruginosa* PA14 to introduce the mutation *pilT-T216R* | This study |
| pMQ30-*pilT-H222A* | For performing allelic exchange at the native locus of the *pilT* gene in *P. aeruginosa* PA14 to introduce the mutation *pilT-H222A* | This study |
| pMQ30-*pilT-H229A* | For performing allelic exchange at the native locus of the *pilT* gene in *P. aeruginosa* PA14 to introduce the mutation *pilT-H22A* | This study |
| pMQ30-*pilT-K136A* | For performing allelic exchange at the native locus of the *pilT* gene in *P. aeruginosa* PA14 to introduce the mutation *pilT-K136A* | This study |

**Supplementary Table S3. Primers used in this study.**

| Primer name | Primer sequence (5’->3’) |
|---|---|
| D17G_pilT_F | CAAACAGGGCGCTTCGGGCCTGCACCTCTCCGCCGGC |
| D17G_pilT_R | GCCGGCGGAGAGGTGCAGGCCCGAAGCGCCCTGTTTG |
| K45E_pilT_F | CCACCGCTGGAACACGAGCAGGTGCATGCGC |
| K45E_pilT_R | GCGCATGCACCTGCTCGTGTTCCAGCGGTGG |
| H48R_pilT_F | GGAACACAAGCAGGTGCGTGCGCTGATCTACGACATC |
| H48R_pilT_R | GATGTCGTAGATCAGCGCACGCACCTGCTTGTGTTCCAG |
| E65K_pilT_F | GCAGCGCAAGGACTTCGAGAAATTCCTCGAGACCGACTTCTCC |
| E65K_pilT_R | GGAGAAGTCGGTCTCGAGGAATTTCTCGAAGTCCTTGCGCTGC |
| N87S_pilT_F | CGGGTCAACGCCTTCAGCCAGAACCGTGGCGC |
| N87S_pilT_R | GCGCCACGGTTCTGGCTGAAGGCGTTGACCCG |
| N87D_pilT_F | CGGGTCAACGCCTTCGACCAGAACCGTGGCGC |
| N87D_pilT_R | GCGCCACGGTTCTGGTCGAAGGCGTTGACCCG |
| H222A_pilT_F | CGCGGCGGAGACCGGCGCCCTGGTATTCGGCACCC |
| H222A_pilT_R | GGGTGCCGAATACCAGGGCGCCGGTCTCCGCCGCG |
| M301T_pilT_F | CGAGGACAAGGTCGCGCAGACGTATTCGGCGATCCAGACC |
| M301T_pilT_R | GGTCTGGATCGCCGAATACGTCTGCGCGACCTTGTCCTCG |
| R331H_pilT_F | GGGCCTGATCAGCCACGAGAATGCCCGCGAGAAGG |
| R331H_pilT_R | CCTTCTCGCGGGCATTCTCGTGGCTGATCAGGCCC |
| YFP_K213E_F | gcaaactgtctaaagacccgaacgaaaaacgtgaccacatgg |
| YFP_K213E_R | ccatgtggtcacgtttttcgttcgggtctttagacagtttgc |
| LBB2H_pT_5' | GCGTCTAGAGATGGATATTACCGAGCTGCTCG |
| LBB2H_pT_3' | GCGGGTACCTCAGAAGTTTTCCGGGATCTTC |
| LBB2H_pU_5' | GCGTCTAGAGATGGAATTCGAAAAGCTGCTGC |
| LBB2H_pU_3' | GCGGGTACCGCTGGCCTACTGAAGACGGT |
| LBB2H_pC_5' | TATATAACTCTAGAGATGGCGGACAAAGCGTTAAAAACCAG |
| LBB2H_pC_3' | TATATATGGAATTCGTTATCCGACGACGTTGCCGA |
| LBB2H_pB_5' | TATATATATCTAGAGATGAACGACAGCATCCAACTG |
| LBB2H_pB_3' | TATATATAGAATTCGTTAATCCTTGGTCACGCGGTT |
| K58A_pilT_F | gatctaCGACATCATGAACGACgcGCAGCGCAAGGACTTCGAGGAATTCC |
| K58A_pilT_R | GGAATTCCTCGAAGTCCTTGCGCTGCgcGTCGTTCATGATGTCGTAGATC |
| H229A_pilT_F | cctGGTATTCGGCACCCTGGCCACCACCTCGGCGGCGAAGACC |
| H229A_pilT_R | GGTCTTCGCCGCCGAGGTGGTggcCAGGGTGCCGAATACCAGG |
| 30pT_1 | CGAATTCGAGCTCGGTACCCCCACGGCCTCGGCGTTGGAC |
| 30pT_6 | GTCGACTCTAGAGGATCCCCCGAACACAGCACCCTGCAACTGGAAACC |
| 30pT_7 | GTCGACTCTAGAGGATCCCCCCAGACCAACTCGACCCGCC |

| | |
|---|---|
| 30pT_2COR | GAGCAGCTCGGTAATATCCAT |
| 30pT_3COR | ATGGATATTACCGAGCTGCTC |
| 30pT_4COR | GAAGTTTTCCGGGATCTTCGC |
| 30pT_5COR | GCGAAGATCCCGGAAAACTTC |
| pilT_D31K_F_COR | CCGCCCATGATCCGGGTGAAGGGCGATGTACGCCGGATCAACCTGCC |
| pilT_D31K_R_COR | GGCAGGTTGATCCGGCGTACATCGCCCTTCACCCGGATCATGGGCGG |
| 72RBS_pilT5' | gGTACCgaaggagatatacatATGGATATTACCGAGCTGCTCGC |
| 72RBS_pilT3' | ccaaaacagccaAGCTTTCAGAAGTTTTCCGGGATCTTCGCC |
| 18C_pilC_XbaI_5' | TATATAACTCTAGAGatggcggacaaagcgttaaagacc |
| 18C_pilC_EcoRI_3' | TATATATGGAATTCGTTAcacaacggaacccagttggaagatcg |
| 72toVLT31_Gib1_5' | CAGAATTCGAGCTCGGTACCctactgtttctccatacccgttttttg |
| 72toVLT31_Gib2_5' | ACACAGGAAACAGAATTCGGctactgtttctccatacccgttttttg |
| PaQaRpoD_CTX1_A | ctagaactagtggatcccccctaataaccaggcatcaaataaaacgaaaggctc |
| PaQaRpoD_CTX1_B2 | atatcgaattcctgcagccgcggccgcaaaaggaaaagatc |
| 72rbs_pT_5' | TcgagctcggtacccCAGGAGGAATTTTCCATGGATATTACCGAGCTGCTCGC |
| 72rbs_pT_3' | CgactctagaggatcccctcaGAAGTTTTCCGGGATCTTCgc |
| T18C_fimL_5' | CGACTCTAGAGGATCCCCGGatggtcacaggagccacgtc |
| T18C_fimL_3' | AATTCGAGCTCGGTACCCTggcggccaccggcag |
| T18C_pilK_5' | CGACTCTAGAGGATCCCCGGatgcaggcgaacggcgtc |
| T18C_pilK_3' | AATTCGAGCTCGGTACCCTtgtgcctgagtaccccttacg |
| T18C_chpB_5' | CGACTCTAGAGGATCCCCGGatgagtgagcgcgccac |
| T18C_chpB_3' | AATTCGAGCTCGGTACCCTtgtttcgactcctgtcggcg |
| T18C_fimW_5' | CGACTCTAGAGGATCCCCGGatggaaaaccagagcccccac |
| T18C_fimW_3' | AATTCGAGCTCGGTACCCTcagagacttccagagcgagtcaaaatc |
| VLT31rbs_pilU_5' | TTCGAGCTCGGTACCCCAGGAGGAATTTTCCatggaattcgaaaagctgctgc |
| VLT31rbs_pilU_3' | CGACTCTAGAGGATCCCCtcagcggaagcgccg |
| T18C_pilG_5' | CGACTCTAGAGGATCCCCGGatggaacagcaatccgacggt |
| T18C_pilG_3' | AATTCGAGCTCGGTACCCTggaaacggcgtccaccg |
| T18C_pilH_5' | CGACTCTAGAGGATCCCCGGatggctcgtattttgattgttgatgact |
| T18C_pilH_3' | AATTCGAGCTCGGTACCCTgcccgccagcaccg |
| T18C_pilC_L_5' | CGACTCTAGAGGATCCCCGGatggcggacaaagcgttaaagac |
| T18C_pilC_S_5' | CGACTCTAGAGGATCCCCGGatgctggtgaaggctcaactg |
| T18C_pilC_3' | AATTCGAGCTCGGTACCCTcacaacggaacccagttggaag |
| T18C_pilU_5' | CGACTCTAGAGGATCCCCGGatggaattcgaaaagctgctgcg |
| T18C_pilU_3' | AATTCGAGCTCGGTACCCTgcggaagcgccgg |

| | |
|---|---|
| T18C_pilI_5' | CGACTCTAGAGGATCCCCGGatgtcggacgttcagaccc |
| T18C_pilI_3' | AATTCGAGCTCGGTACCCTtacggcgacgtcgagga |
| T18C_chpC_5' | CGACTCTAGAGGATCCCCGGatgaaccaggccgtgatcgag |
| T18C_chpC_3' | AATTCGAGCTCGGTACCCTgatcaggccggcgtcg |
| T18C_chpD_5' | CGACTCTAGAGGATCCCCGGatggccggcctgcaac |
| T18C_chpD_3' | AATTCGAGCTCGGTACCCTgccgcgcacagcg |
| T18C_chpE_5' | CGACTCTAGAGGATCCCCGGatgctcgccatcttcctcg |
| T18C_chpE_3' | AATTCGAGCTCGGTACCCTcagcccgcgcagc |
| QC_pilT_R123D_5' | CGTGTTTCAGACGTCCCGgaCGGGCTGGTACTGGTCACCG |
| QC_pilT_R123D_3' | CGGTGACCAGTACCAGCCCGtcCGGGACGTCTGAAACACG |
| QC_pilT_T216R_5' | CGCCTGGCCCTGAgaGCGGCGGAGACCGGCC |
| QC_pilT_T216R_3' | GGCCGGTCTCCGCCGCtcTCAGGGCCAGGCG |

**References.**
